# Revisiting the phosphorite deposit of Fontanarejo (central Spain): new window into the early Cambrian evolution of sponges and into the microbial origin of phosphorites

**DOI:** 10.1101/2020.12.13.422563

**Authors:** Joachim Reitner, Cui Luo, Pablo Suarez-Gonzales, Jan-Peter Duda

**Affiliations:** Department of Geobiology, Centre of Geosciences of the University of Göttingen Goldschmidtstraße 3; Academy of Science and Humanities, Theater Str. 7, 37077 Göttingen, Germany; State Key Laboratory of Palaeobiology and Stratigraphy, Nanjing Institute of Geology and Palaeontology and Center for Excellence in Life and Paleoenvironment, Chinese Academy of Sciences, 39 East Beijing Road, Nanjing 210008, China; Departamento de Geodinámica, Estratigrafía y Paleontología, Universidad Complutense de Madrid, C/ José Antonio Novais 12, 28040 Madrid, Spain; Sedimentology & Organic Geochemistry Group, Department of Geosciences, Eberhard-Karls-University Tübingen, Schnarrenbergstraße 94-96, 72076 Tuebingen Germany

**Keywords:** Lower Cambrian, Porifera, Hexactinellida, Demospongiae, microbialites

## Abstract

Fossils within early Cambrian phosphorites worldwide are often well preserved due to early diagenetic permineralization. Here, we examine the fossil record contained within phosphorites of the Lower Cambrian Pusa Formation (late Fortunian to Cambrian Stage 2) in Fontanarejo, central Spain. The sedimentology and age of these phosphorites have been controversial and are here reviewed and discussed, providing also a updated geological map. The Pusa Formation is composed of fine clastic sediments that are partly turbiditic, with channels of quartz-rich conglomerates and abundant phosphorites in the upper part of the succession. The microfacies and mineralogy of these channel deposits are studied here for the first time in detail, showing that they are mainly composed of subspherical apatite clasts, with minor mudstone intraclasts, quartzite and mica grains. Numerous sponge spicules, as well as entirely preserved hexactinellid sponges and demosponges, were collected within these phosphorites and likely represent stem groups. In addition to sponges, other fossils, such as small shelly fossils (SSF) of the mollusk *Anabarella* sp., were found. The phosphorites exhibit multiple evidence of intense microbial activity, including diverse fabrics (phosphatic oncoidal-like microbialites, thrombolites, stromatolites, and cements) and abundant fossils of filamentous microbes that strongly resemble sulfur oxidizing bacteria. Our findings strongly suggest that microbial processes mediated the rapid formation of most of the Fontanarejo apatite, probably accounting for the exceptional preservation of fragile fossils such as sponge skeletons. The apparent presence of taxonomically diverse hexactinellid and demosponge communities by the lowermost Cambrian further corroborates a Precambrian origin of the phylum Porifera.

## 1. Introduction

The poor early fossil record makes investigations of sponge evolution difficult. Discounting the questionable Ediacaran sponge-like fossils (e.g. Gehling & Rigby, 1996; Brasier *et al*. 1997; Li *et al*. 1998; Reitner & Wörheide, 2002; Maloof *et al*. 2010; Antcliffe *et al*. 2014), the earliest appearance of sponge spicules has been dated to ca. 535 Ma (Guo *et al*. 2008; Chang *et al*. 2017). Problematically, however, disarticulated spicules are of little taxonomic value. To date, articulated, well-preserved sponge fossils in Burgess-Shale-type preservation and deep-water black shales have only been reported as after the beginning of Cambrian Stage 3 (e.g. Steiner *et al*. 1993, Luo *et al*. 2020). Furthermore, Luo & Reitner (2019) described three-dimensionally preserved sponge skeletal frames from the basal Niutitang Formation (Stage 2) near Sancha in China that have illuminated the nature of stem-group hexactinellids.

Here, new occurrences of sponge fossils of no later than the Cambrian Stage 2 are described from phosphorite deposits near the village of Fontanarejo in central Spain (Fig. 1). The Fontanarejo phosphorites belong to the Pusa Formation and were targets of mining exploration (IGME *et al*. 1984-1987). Perconig *et al*. (1983, 1986) provided the first summary of the formation’s stratigraphy, sedimentology, and fossil content (particularly sponge spicules). While these early studies considered the Fontanarejo phosphorites to be of Ediacaran age, Jensen *et al*. (2010) suggested a Lower Cambrian age (Fortunian) for the Pusa Formation. Reitner *et al*. (2012) provided the first overview of the different sponge types contained within the Fontanarejo outcrops, highlighting their stratigraphic value. The aim of this paper is to describe in more detail the different types of sponge remains found in the Fontanarejo phosphorites and settle them to the current taxonomic frame. In addition to sponges, other fossils recovered from the phosphorites, including small shelly fossils (SSF) and microbial fossils, were studied. A further goal of the paper is to tackle the microbial impact of an early diagenetic apatite formation to explain the exceptional preservation of fossils.

**Fig. 1.**
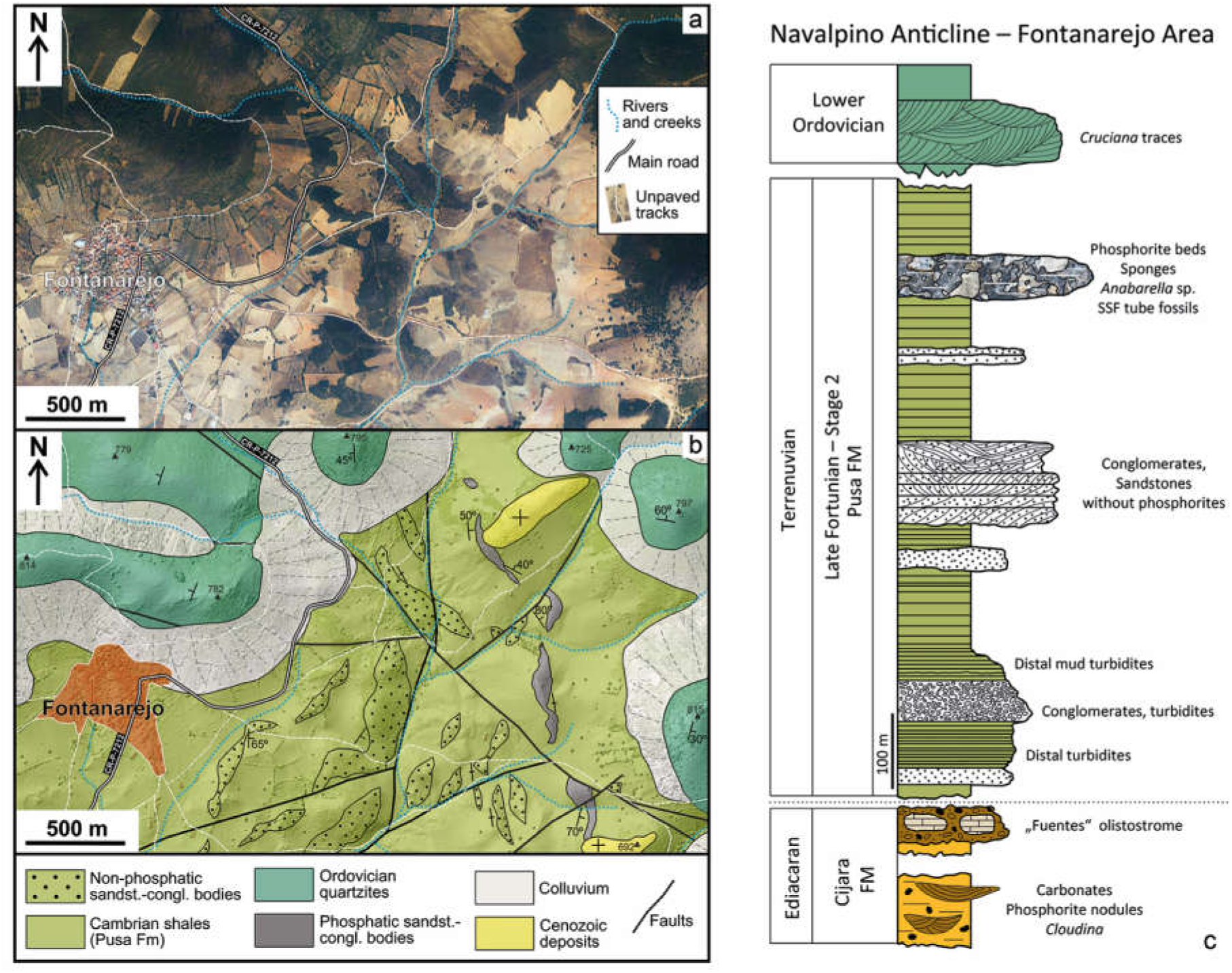
(a) Latest available orthophotograph of the Fontanarejo area (obtained from the Spanish Geographical Survey, IGN) showing the current outcrop conditions, dominated by vegetation and agricultural lands. (b) Geological map of the Fontanarejo area, same extension as in (a), at the western end of the Navalpino Anticline. Map is drawn over a LIDAR digital elevation model (obtained from the Spanish Geographical Survey, IGN) and based on our own fieldwork and on the previous geological maps of the area. (c) Reconstructed stratigraphic section of the Fontanarejo area, based on Perconig *et al*. (1986), Picart Boira (1988), IGME (1984–1987), and own fieldwork. The Ediacaran Cijara Formation and the so-called ‘Fuentes’ olistostromes, which roughly represent the Precambrian-Cambrian boundary, are not exposed in this area. Data used in this reconstruction are from Álvaro *et al*. (2016) and Jensen *et al*. (2010). This section represents the stratigraphic base, which is exposed in the Valdelacasa Anticline west of Fontanarejo.

Assuming that animal life had a long evolutionary history before the Cambrian, paleontologists have been tracing the earliest fossil evidence of sponges for many decades. Putative Precambrian spicule-like structures have been reported from analyses of thin sections (e.g. Brasier *et al*. 1997; Li *et al*. 1998; Reitner & Wörheide, 2002), and macroscopic Ediacaran fossils (e.g. Gehling & Rigby, 1996; Serezhnikova & Ivantsov, 2007; Clites *et al*. 2012). Furthermore, enigmatic structures observed in various Neoproterozoic successions were interpreted to be poriferan (e.g. Maloof *et al*. 2010; Brain *et al*. 2012; Wallace *et al*. 2014). These interpretations have mostly been discounted by later studies (e.g. Antcliffe *et al*. 2014; Muscente *et al*. 2015; Cunningham *et al*. 2017; Luo *et al*. 2017). The most plausible alternative evidence for Precambrian sponges, that is molecular fossils (e.g. Love *et al*. 2009; Zumberge *et al*. 2018) and exceptionally preserved fossils from the Weng’an Biota (Yin *et al*. 2015), are also still under debate (Botting & Muir, 2018; Nettersheim *et al*. 2019).

Despite the possibility that the earliest sponges were too small or labile to be fossilized, or that they looked very different from their Phanerozoic descendants, the earliest indisputable sponge fossils are disarticulated spicules recognized from the Fortunian (538.8–529.0 Ma, Peng *et al*. 2020). These fossil spicules include pentactins and hexactins macerated from the *Protoherzina* biozone of the Yangjiaping Formation of Hunan, China (Ding & Qian, 1988), and stauractins and pentactins in the same biozone of the Soltanieh Formation in North Iran (Antcliffe *et al*. 2014). The spicules from Iran occur right above the Basal Cambrian Carbon Isotope Excursion (BACE). Probably even earlier occurrences of spicules were reported from the Yanjiahe Formation of Yangtze Gorges which were correlated to around the nadir of BASE, although the referred geochemical profile were not directly obtained from the fossil-containing outcrop (Chang *et al*. 2017, 2019).

Taxonomically more informative sponge fossils (i.e., those preserved with articulated skeletal frames) are mostly known from the fossil Lagerstätten shales since slightly before the Cambrian Stage 3. Famous examples include the black shales of the Niutitang and Hetang Formations in South China (Steiner *et al*. 1993; Xiao *et al*. 2005), the Chengjiang Biota (e.g. Chen *et al*. 1989, 1990; Rigby & Hou 1995), the Sirius Passet (Botting & Peel, 2016), and the Burgess Shale fossil Lagerstätte (e.g. Rigby 1986; Rigby & Collins 2004). The recently described three-dimensionally preserved hexactinellid sponge fossils and diverse disarticulated spicules from the cherty phosphorites at the basal Niutitang Formation of Hunan, China, were probably no younger than 521 Ma, thus providing a valuable window for studying sponge paleobiodiversity, paleoecology, and evolution preceding the Cambrian Series 2 (Luo & Reitner, 2019).

Few sponge fossils from the Fontanarejo phosphorites were also three-dimensionally preserved and have been briefly described in Perconig *et al*. (1986), Reitner *et al*. (2012), and Reitner & Luo (2019). This Fontanarejo fossil Lagerstätte is partly to the Niutitang phosphorites in age—though slightly older—based on the more recent discovery of *Anabarella* and improved sedimentological correlation (Álvaro *et al*. 2019; this study). It therefore provides a unique and valuabe window into early metazoan diversity, particularly allowing for the reconstruction of the major sponge branches.

Generally, the classification used here follows the traditional concept of the encyclopodia *Systema Porifera: A Guide to the Classification of Sponges* (Hooper & Van Soest, 2002). In addition, we integrate findings from the most recent phylogenetic models provided by Erpenbeck & Wörheide (2007), Dohrmann *et al*. (2008), Wörheide *et al*. (2012), and Redmond *et al*. (2013). Combining fossil and modern concepts always involves taxonomic compromise.

## 2. Material and methods

Over the last two decades, we conducted several field campaigns to the phosphorite deposits in the Fontanarejo area, with the aim of mapping the outcrop and collecting samples. In the field, we collected data for preparing a new geological map of the area, using previous maps as a starting point (Perconig *et al*. 1983, 1986; IGME *et al*. 1984–1987; MAYASA *et al*. 1987–1990, 1990–1993; Monteserín López *et al*. 1989; López Díaz, 1995) (Fig. 1). The background geographic information used was the latest available topographic maps, LIDAR digital elevation models, and orthophotos obtained from the Spanish Geographical Survey (IGN). The geographic data were combined and managed using QGIS software, and the geological map was drawn using CorelDraw.

We collected more than 200 rock samples in the field and prepared 120 large thin sections (10 x 5 cm, 5 x 5 cm, and 15 x 10 cm) from them. The thin sections are archived in the museum collection of the faculty of Geoscience and Geography in Göttingen. Thin section analyses were conducted using a Zeiss SteREO Discovery.V12 stereo microscope and a Zeiss AxioImager Z1 microscope. Both microscopes were connected to a Zeiss AxioCam MRc camera. A fluorescence microscope equipped with an ebx 75 isolated power unit was mounted on a Zeiss AxioImager Z1 microscope. The instrument was fitted with a Zeiss A 10 Alexa Fluor 488 filter, with an excitation wavelength of BP 450–490 nm and an emission wavelength of BP 515–565 nm. For cathodoluminescence (CL) microscopy, a Cambridge Instrument Citl CCL 8200 Mk3A cold-cathode system (operating voltage of c.15 kV; electric current of c. 250–300 μA) linked to a Zeiss Axiolab microscope and AxioCam 703 camera was used. Field emission scanning electron microscopy (Fe-SEM) was performed with a Carl Zeiss LEO 1530 Gemini system. The instrument was coupled to an Oxford Instruments INCA x-act energy dispersive X-ray spectrometry (EDX) detector, to additionally obtain EDX single spectra and elemental maps.

Phosphorite mineralogy was analyzed using micro x-ray fluorescence (μ-XRF); a Bruker M4 Tornado equipped with a XFlash 430 Silicon Drift Detector obtained elemental distributions within the thin sections. Analyses were carried out at a spatial resolution of 70 µm. Each pixel covers 30 ms. Measurements were performed at 50 kV and 400 µA at a chamber pressure of 20 mbar. Additionally Raman spectroscopy was used for point measurements and area mapping. Raman spectra were collected using a WITec alpha300 R fiber-coupled ultra-high throughput spectrometer. Before analysis, the system was calibrated using an integrated light source. The experimental setup included a 405 nm laser (chosen to reduce fluorescence effects), 10 mW laser power, and 50x and 100x long working distance objectives with a numerical aperture of 0.55 and a grating of 1200 g mm^−1^. This setup had a spectral resolution of 2.6 cm^−1^. The spectrometer was centered at 1530 cm^−1^, covering a spectral range from 122 cm^−1^ to 2759 cm^−1^. Each spectrum was collected by two accumulations, with an acquisition time of 5 s. Raman spectra were processed with the WITec project software. The background was subtracted using a rounded shape, and band positions were determined by fitting a Lorentz function. Both analytical facilities (µ-XRF and Raman spectroscopy) are located at the Geosciences Center of the University of Göttingen.

## 3. Regional geology and stratigraphy

### 3.a. Regional Geological Setting

The studied deposits crop out near the village of Fontanarejo (Ciudad Real province, central Spain). Geographically they are located at the southern margin of the Toledo Mountains, and geologically they are part of the southeast area of the Central Iberian Zone (CIZ) of the Iberian Variscan Massif. This area of the CIZ consists of large-scale, NW-SE trending upright folds, topographically highlighted by the relief of the Ordovician quartzites. The folds include wide anticlines containing pre-Ordovician materials and narrower synclines containing Ordovician-Devonian deposits (Díez Balda *et al*. 1990; Vidal *et al*. 1994; Liñán *et al*. 2002; Valladarez *et al*. 2002; Martínez Catalán *et al*. 2004; Álvaro *et al*. 2019). The Fontanarejo area represents the northeastern border of the Navalpino Anticline, which accordingly exhibits a core of folded and deformed Neoproterozoic and Cambrian rocks unconformably overlain by Ordovician quartzites (San José, 1984; López Díaz, 1994, 1995; Gutiérrez-Marco *et al*. 2017) (Figs. 1b, c, 2). The studied phosphate-rich deposits are part of the Cambrian succession that directly underlies the Ordovician quartzites of the northeastern margin of the Navalpino Anticline (cf. Perconig *et al*. 1983, 1986; Picart Boira, 1988) (Figs. 1b, c, 2).

### 3.b. Stratigraphy and Age

The stratigraphic terminology and age of the pre-Ordovician deposits of the CIZ have been controversial for several decades (see discussions in San José *et al*. 1990; Vidal *et al*. 1994; Rodríguez Alonso *et al*. 2004; Jensen *et al*. 2010; Álvaro *et al*. 2019). Typically, these deposits have been divided into three large lithostratigraphic units, with varying names and boundaries. The Fontanarejo phosphorites belong to the youngest of these groups, termed “Pusian” (San José *et al*. 1990), “Valdelacasa Group” (Monteserín López *et al*. 1989), or “Río Huso Group” (Vidal *et al*. 1994). Within this group, the studied deposits belong to a shale-dominated unit called the Pusa Formation, whose composition and boundaries have also been studied previously (e.g. San José *et al*. 1990; Vidal *et al*. 1994; Álvaro *et al*. 2019).

From a sedimentological point of view, the Pusa Formation in our study area has six sequences or subunits (San José, 1984; Gabaldón López *et al*. 1989), with the Fontanarejo phosphorites being part of subunit 5. San José (1984) even defined them as own member within the unit (”Fontanarejo Member”). From a cartographic point of view, the Pusa Formation has only two (Monteserín López *et al*. 1989) or three (San José, 1984; López Díaz, 1994) mappable subunits. The latest stratigraphic review (Álvaro *et al*. 2019) considers three mappable lithostratigraphic members, with the Fontanarejo phosphorites part of the middle, conglomerate-rich member (Figs. 1c, 2). According to this recent stratigraphic framework, the Pusa Formation is as thick as 3500 m in places and is dominated by shales interbedded with breccia, conglomerates, and sandstones, as well as local carbonate and phosphorite seams (Álvaro *et al*. 2019).

**Fig. 2.**
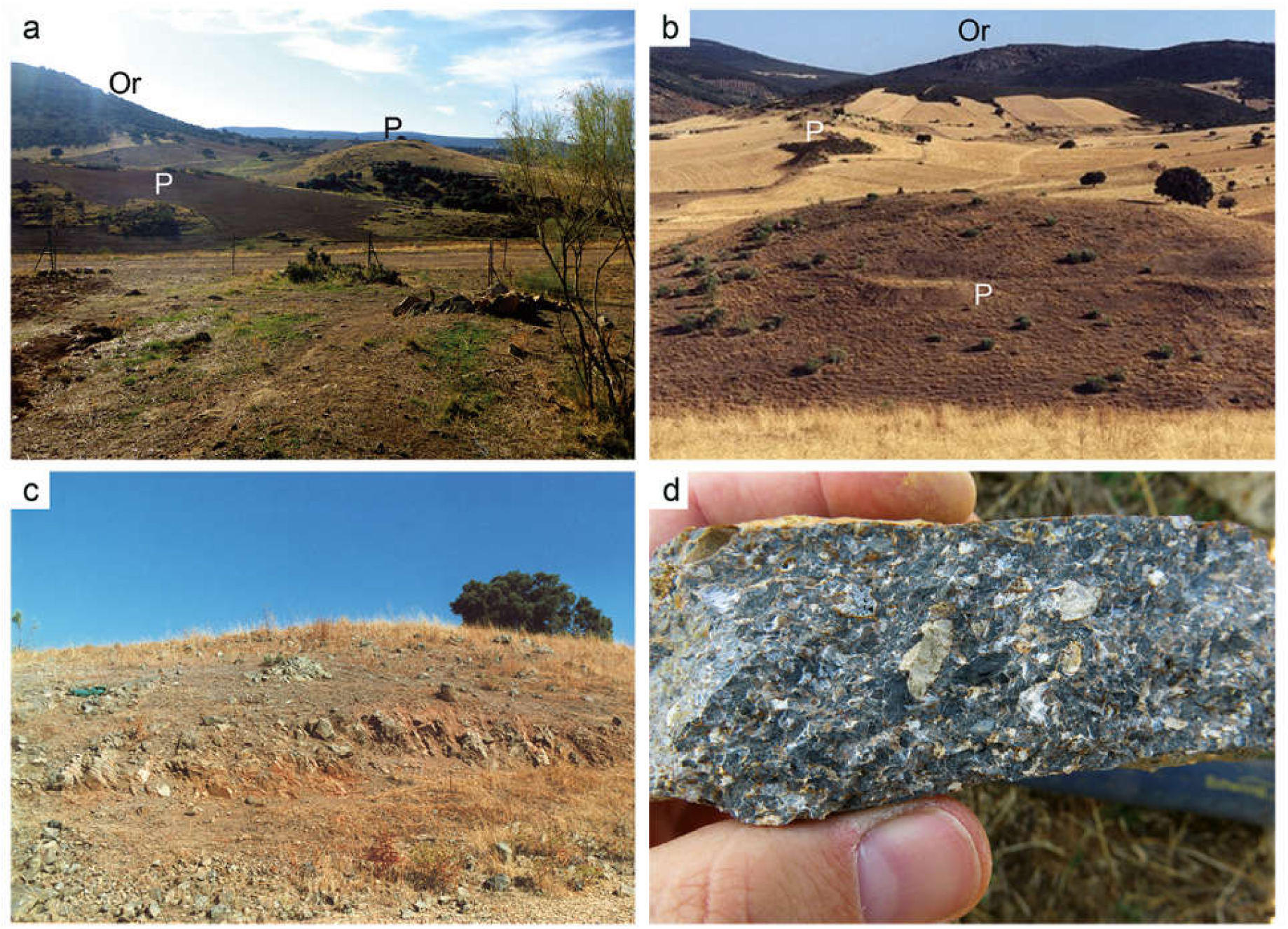
(a) Landscape view to the south with hills showing phosphatic sediments (P). Forest-rich areas represent Ordovician quartzites (Or). (b) Landscape view to the north, with hills showing phosphatic sediments (P). (c) Bedded phosphatic sediments near elevation point 692 m in the southeast corner of the geological map. Picture obtained in 2002 – today this area is planted with trees. The beds show an inclination to the east of 70°. (d) Detailed picture of coarse phosphatic sediment from Fig. 2c.

The age of the Pusa Formation remains uncertain, but most interpretations agree that it includes deposits from the latest Neoproterozoic and earliest Cambrian (Brasier *et al*. 1979; Perconig *et al*. 1983, 1986; San José, 1984; Monteserín López *et al*. 1989; Vidal *et al*. 1994; Jensen *et al*. 2010). However, a zircon age of 533 ± 17 Ma from the middle part of its lower member (Talavera *et al*. 2012), and the discovery of *Anabarella* cf. *plana* fossils, have led the Pusa Formation to be assigned exclusively to the Lower Cambrian, mainly to the Terreneuvian, with only its uppermost deposits possibly being Cambrian Series 2 in age (Álvaro *et al*. 2019).

Notably, we also found *Anabarella*-like fossils within the Fontanarejo phosphorite deposits. This finding aligns with occurrences of *Anabarella* cf. *plana* in stratigraphically equivalent sedimentary rocks from the Alcudia anticline close to Ciudad Real in central Spain (i.e., “Conglomerados de San Lorenzo”: Pieren & García-Hidalgo, 1999; Vidal *et al*. 1999; Pieren Pidal, 2000; Álvaro *et al*. 2019). Given this stratigraphic framework, the phosphorite deposits of Fontanarejo we studied are likely late Fortunian – base of Stage 2 in age. Figure 1c summarizes the main sedimentary structures and the stratigraphy of the Fontanarejo area.

## 4. Outcrop Description and Sedimentology

The landscape of the Fontanarejo area is mainly controlled by the weathering susceptibility of shales from the Pusa Formation that underlie the main agriculture fields (Figs. 2). Interbedded with the shales are more weathering-resistant sandstone-conglomerate beds that form the small hills of the landscape. The phosphorites we studied crop out ∼2 km east of the village of Fontanarejo and are embedded in the youngest series of the sandstone-conglomerate beds (Perconig *et al*. 1983, 1986; Picart Boira, 1988) (Figs. 1b, c; 2, 3). These phosphate deposits were first identified in the course of extensive field campaigns conducted by the Spanish Geological Survey (IGME) during the 1980s (IGME *et al*. 1984–1987; MAYASA *et al*. 1987–1990, 1990–1993). Since then, the Fontanarejo outcrops have been significantly degraded due to agricultural activities (Fig. 2). This makes it almost impossible to find *in situ* outcrops larger than a couple square meters, and those are significantly weathered. A further problem is that different phosphate beds cannot be directly correlated due to fault displacement.

Given the current poor outcrop conditions, it is difficult to accurately assess the geometry and macroscopic features of the sandstone-conglomerate beds in greater detail. However, they are clearly discontinuous and locally seem to pinch out laterally. Furthermore, they appear to be 100–500 m wide and 20–70 m thick. All these characteristics are consistent with the previous descriptions of the surface and subsurface (IGME *et al*. 1984–1987; MAYASA *et al*. 1987–1990). Notably, these studies also reported that some of the beds display erosional bases and flat to undulated tops and internally include fining-upward, cross-bedded sediments (Perconig *et al*. 1983, 1986; IGME *et al*. 1984–1987; MAYASA *et al*. 1987–1990, 1990–1993; Picart Boira, 1988; Álvaro *et al*. 2016). We found these features only to be present in sandstone-conglomerate beds that do not contain significant number of phosphate pebbles (Fig. 3e), while they appear to be absent in the youngest, phosphate-rich beds in the succession (Figs. 2, 4). Regardless of these differences, however, integrated map and field evidence suggest that the phosphate-rich and the phosphate-poor bodies are both channel deposits.

**Fig. 3.**
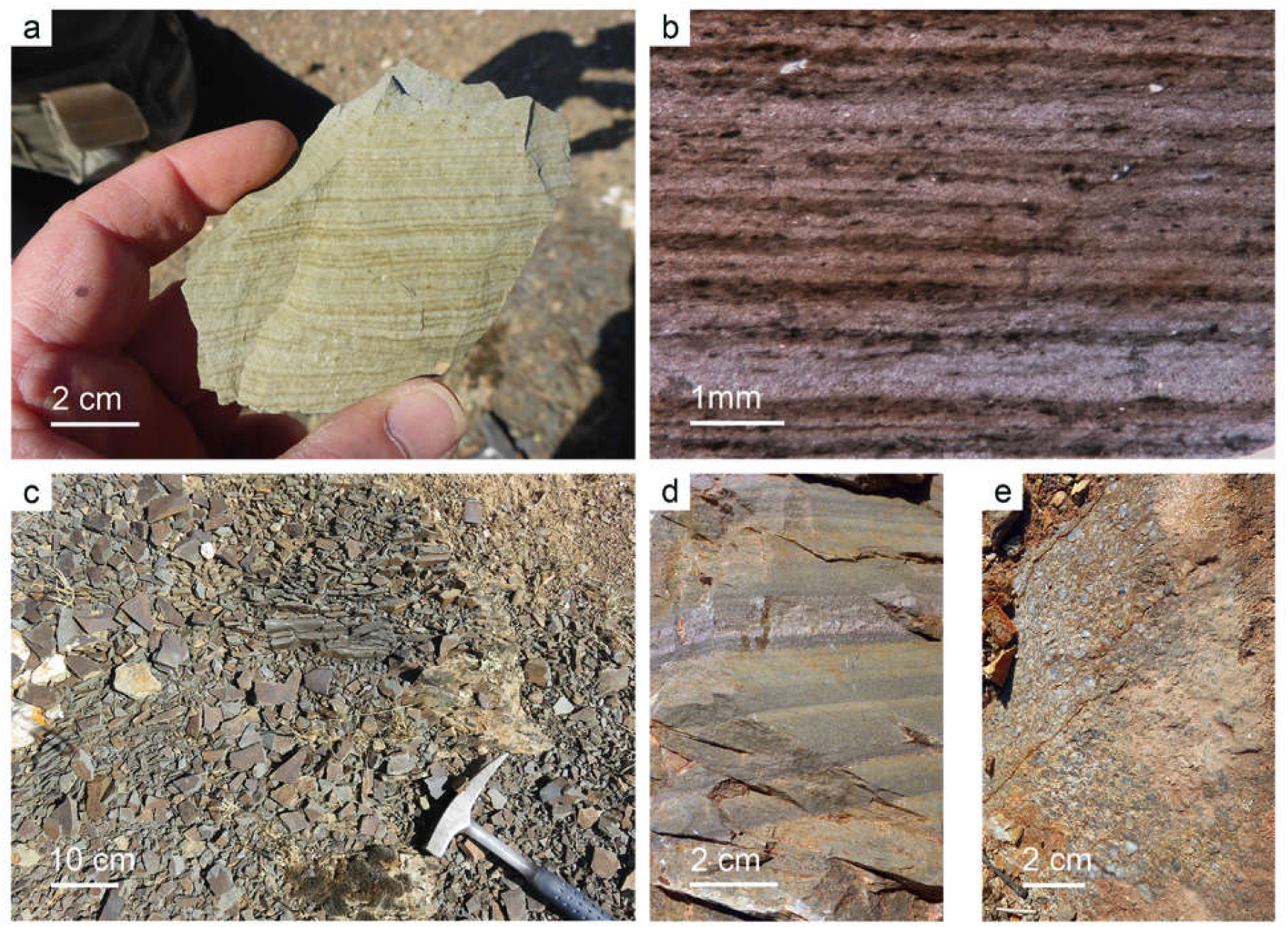
Sedimentary structures of the Pusa Formation. (a) Specimen of Pusa Fm. exhibiting fine lamination of distal turbidites. (b) Thin section micrograph of the panel in Fig. 3a showing bright silt-rich layers overlain by dark mud layers (Bouma Td-e). (c) Gray silt-rich Pusa mudstones. (d) Distal turbidite layers (Bouma Tc-e). (e) Meter-thick channel-filling conglomerate (without phosphorite).

Detailed microfacies analyses of the youngest, phosphate-rich sandstone-conglomerate beds revealed that these deposits exhibit a typical rudstone fabric (Fig. 2d, 4). The phosphate clasts are ∼90% ovoidal to subspherical and display black, gray, and brown colors. The remaining 10% of the fill includes reworked, sub-rounded clasts Pusa shale, often containing angular gray mudstone intraclasts and quartzite grains with minor bits of mica. Iron hydroxide crusts surround many phosphate clasts and are likely remnants of pyrites (remains of framboidal aggregates). Two types of phosphatic sediments are present, one is more weathered and iron hydroxide rich, the other is fresher, with black to gray clasts with less iron hydroxide.

The phosphate clasts consist of apatite and display diverse morphologies. Ovoid-shaped clasts, some millimeter in size (“oncoidal-like microbialites”), are abundant and have distinct nuclei. They are often composed of sponge remains, SSF relics, and mud clasts. Anatase (TiO_2_) is common and often associated with microbial remains, rendering an authigenic origin likely. The oncoidal-like microbialites do not show any clear laminated fabric but instead display cloudy colloform, often thrombolitic structures that are enriched in organic matter. Clay minerals form the nucleic area of the oncoidal-like microbialites (Fig. 4).

**Fig. 4.**
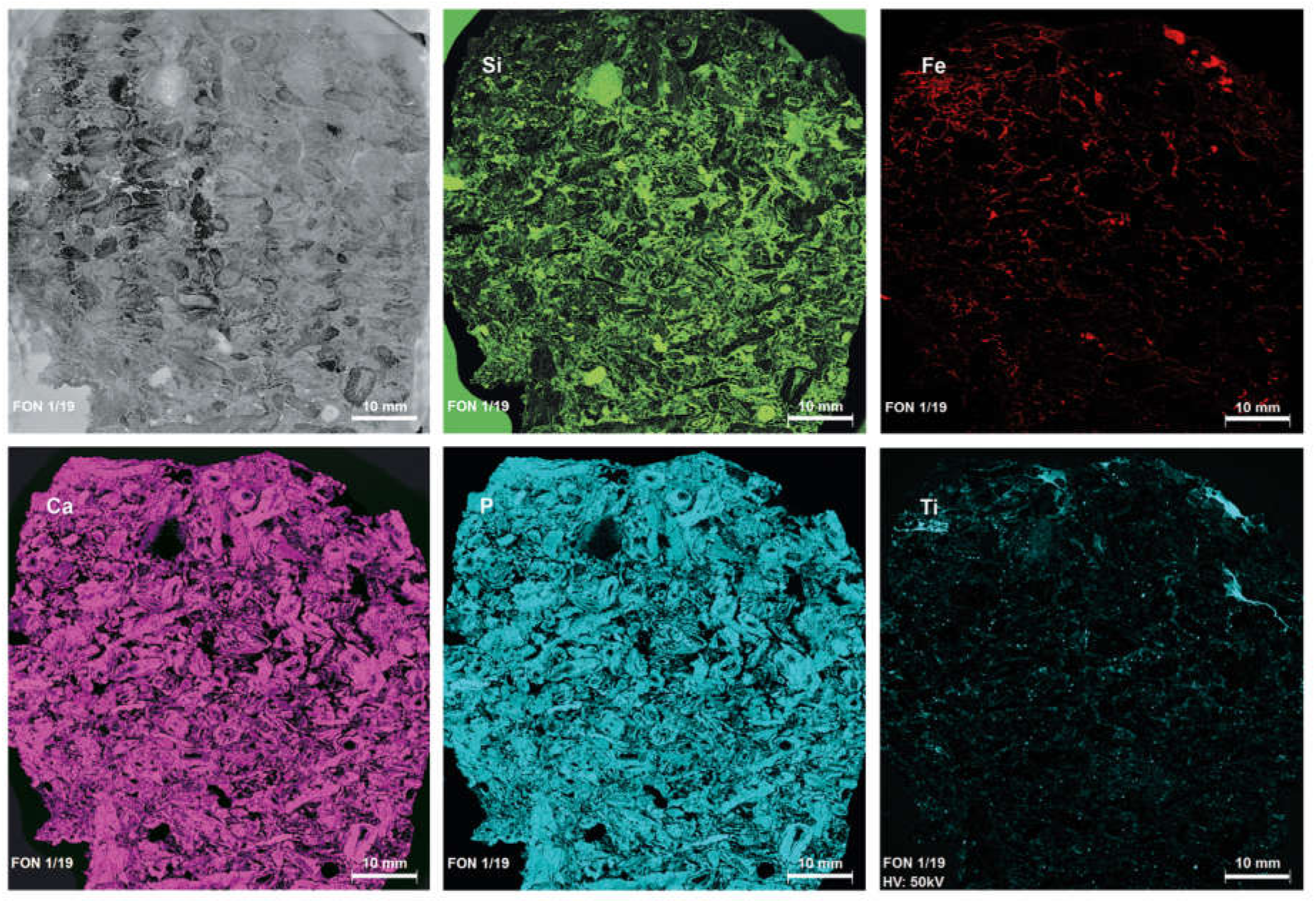
µ-X-Ray fluorescence (-XRF) maps of a thin section of phosphatic sediment (Fon1/19) showing the distribution of major facies-related elements. The rock mainly cemented by silica (Si). Iron (Fe) distribution related with abundant Fe-hydroxide crusts. Fe-hydroxides are probably oxidized phases of former pyrite. Calcium (Ca) and phosphor (P) is most abundant and represent the Ca-P mineral apatite. Intriguing are high amounts of TiO_2_.

In addition to the oncoidal-like microbialites, mm-sized microcrystalline structureless apatite clasts are also common and often include sponge spicules. The microcrystalline apatite components usually show shrinkage patterns and are brown or dark gray in color. The spicule-rich apatite, in contrast, has an irregular shape, is generally honey-yellow in color, and is locally enriched in organic matter. Such micropeloidal textures typically result from the taphonomic mineralization of microbially interspersed sponge tissue (Reitner & Neuweiler, 1995; Reitner & Schumann-Kindel, 1997). For this reason, we interpret these components as permineralized sponge tissue.

Early diagenetic phosphatic cements are quite common. Early prismatic phosphatic cements surround components, spicules, and other components. This early cementation fused original spicule arrangements, on this way greatly facilitating the interpretation of the sponge fossils. Later diagenetic phosphatic cements exhibit larger crystals, and it seems that phosphate minerals replaced siliceous (spicules) and calcareous materials (cf. Perconig *et al*. 1983, 1986; Castaño Prieto, 1987; Picart Boira, 1988; Álvaro *et al*. 2016).

Most studies interpret the Fontanarejo phosphorites (and equivalent deposits in adjacent areas) as allochthonous channel deposits formed in an offshore shelf/platform setting, below wave base level, probably close to the platform margin or to the beginning of the slope (Picart Boira, 1988; Gabaldón López *et al*. 1989; Pieren Pidal, 2000; Álvaro *et al*. 2016). However, Perconig *et al*. (1983, 1986) interpreted the Fontanarejo phosphorites as intertidal and subtidal channel deposits, based on the sorting and rounding of the clasts, the biogenic processes recognized in the rocks, and the local presence of mud cracks. Our findings clearly support the first, more traditional interpretation. This conclusion is based on map and field evidence as well as on microfacies characteristics (see above). For instance, we found no sedimentological evidence for subaerial exposure (e.g. mud cracks). Furthermore, the channel infill is polymictic and poorly sorted, suggesting relatively rapid sedimentation after high-energy episodes that remobilized material from shallower areas as well as material from adjacent zones. The interpretation that some of the Pusa shales are distal mud turbidites interbedded with channels and turbiditic sandstone layers is intriguing (Figs. 1b, 3a, b, d) and consistent with the well-established regional interpretation of the formation as turbidites deposited in the platform-slope transition of a passive margin (San José, 1984; Monteserín López *et al*. 1989; López Díaz, 1994; Vidal *et al*. 1994; Rodríguez Alonso *et al*. 2004; Álvaro *et al*. 2016, 2019).

## 5. Results and discussion: Animal fossils in Fontanarejo phosphorites

### 5.a. Phylogentic status of Hexactinellida Schmidt, 1870

Hexactinellid sponges are characterized by triaxon spicules. Hexactinellids are a monophyletic group, and based on distinct microscleres, two major clades are defined: Hexasterophora and Amphidiscophora (Schulze, 1886). The basic stem group hexactinellid spicule is a regular hexactin (Reitner & Mehl, 1996). How to include the fossil record in the modern molecular phylogenetic tree has been long debated, therefore, it is more convenient to follow the morphological taxonomic criteria errected by Mehl (1996), Mehl-Janussen (1999), and Leys *et al*. (2007) for the early record.

The skeletal architecture of early Paleozoic hexactinellids can be grouped into two major clades, the Reticulosa Reid, 1958 and the “Rossellimorpha” (Mehl, 1996; Mehl-Janussen, 1999), an ill-defined paraphyletic group of early Cambrian hexactinellids (for a more detailed review see Luo & Reitner, 2019). Most of the Fontanarejo articulated hexactinellid sponges exhibit a lyssacine body plan, a primarly non-cemented spicular skeleton. These early lyssacine hexactinellids belong to a widely accepted group that Mehl-Janussen (1999) call “Rossellimorpha A.” Few Fontanarejo hexactinellids display a cemented skeleton resembling a dictyonal body plan, in which the spicules are cemented by biologically controlled secondary silicate cementation. Dictyonal skeletons may have evolved independently and are therefore a polyphyletic charactertistic. In Steiner *et al*. (1993) a first record of a dictyonal hexactinellid from China was described, *Sanshadictya microreticulata* Mehl & Reitner 1993, which also has a an early Cambrian (stage 3) age. To date, the earliest convincing modern dictyonal skeletons were observed in middle Devonian rocks (Mehl, 1996).

### 5.b. Fontanarejo hexactinellid spicule record

Most of the observed hexactinellid spicules are adaptions of simple hexactins (Figs. 5–7). The average size of the spicules is c. 500 µm. Many show remains of an axial canal that was originally filled with a protein-rich organic fiber but now is often filled with iron hydroxide, likely resulting from the diagenetic alteration of primary framboidal pyrite. Some spicules are entirely replaced by iron hydroxide (Figs. 5g–I, 6a, b). Silicious spicules tend to be completely recrystallized and are typically overgrown by the first generation of carbonate cement that was then diagenetically phosphatized. In addition to simple hexactins, differentiated triaxon spicules were observed, such as dermal pentactins (Figs. 5h, 9d, 10a, b), stauractins (Figs. 5i, 8b), and maybe diactins (Figs. 6b, 7c, 9b). Differentiation in mega- and microscleres, as seen in the ball-shaped hexactinellids of the Niutitang Formation basal phosphorites in China (Luo & Reitner, 2019), was not observed.

**Fig. 5.**
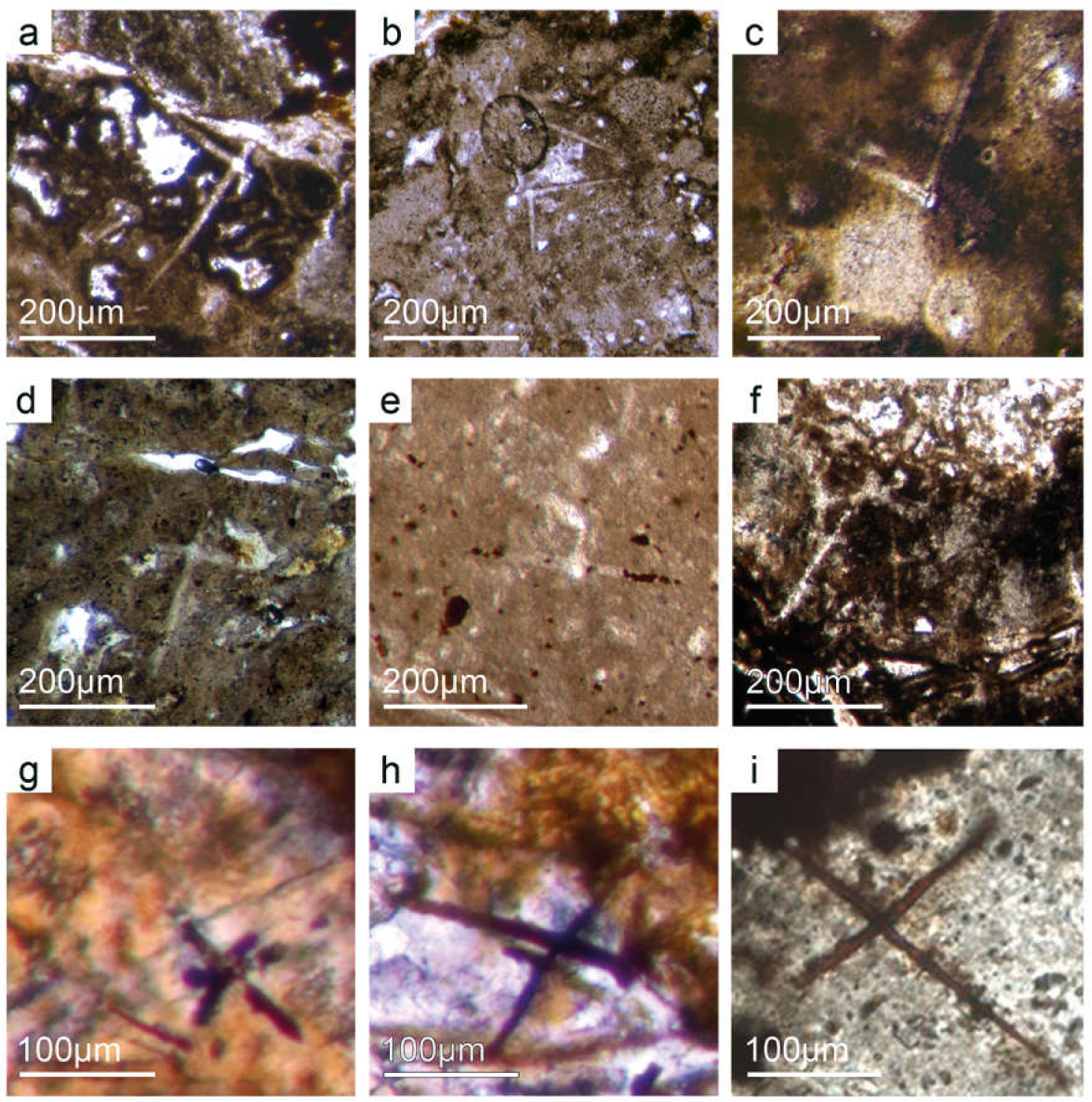
Hexactinellid spicule types. (a–b) Oblique cuts of regular hexactins (Fon1,26). I Hexatin with preserved axial canals (Fon36). (d) A regular hexactin (Fon29). I Stauractin with framboidal pyrite aggregates within spicule rays (Fon19). (f) A regular hexactin (Fon20). (g) A regular hexactine embedded in yellow-brownish transparent phosphate, preserved in Fe-hydroxide that derived from pyrite (Fon27). (h) Pentactin and stauractin embedded in yellow-brownish phosphate, preserved in Fe-hydroxide that derived from pyrite (Fon27). (i) A stauractin preserved in Fe-hydroxide embedded in bright phosphatic material (Fon02).

**Fig. 6.**
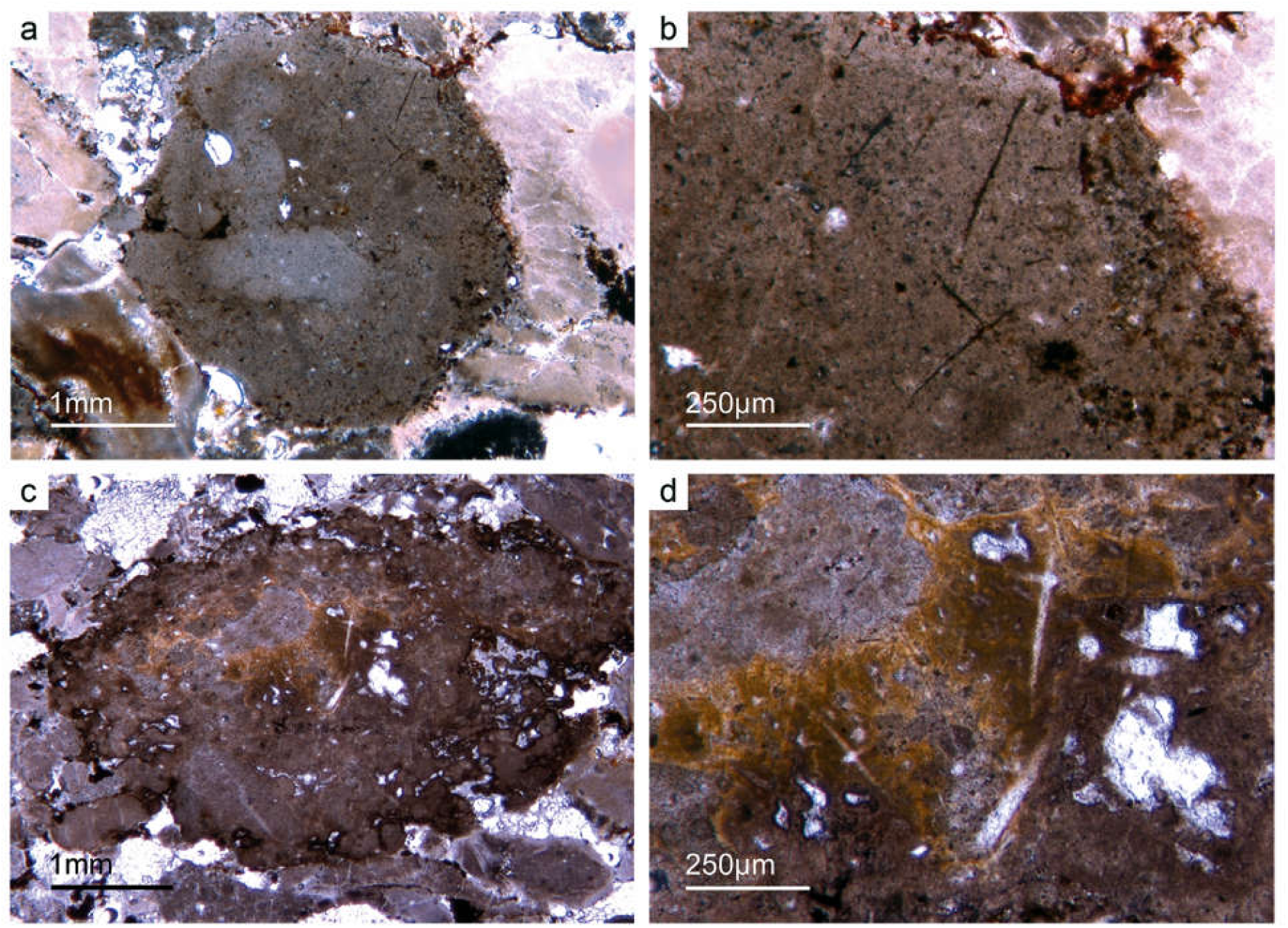
Lyssacine hexactinellid skeletons. (a) A ball-shaped lyssacine hexactinellid preserved in dark-grey phosphate (Fon99). (b) Detail of Fig. 6a showing monaxon and pentactin spicules preserved in Fe-hydroxide – former pyrite. (c) A lyssacine hexactinellid preserved in honey-yellow phosphate (Fon38). (d) Detail of Fig. 6c exhibiting large dermal spicules like pentactins and stauractins. Parenchymal spicules are smaller and hard to see. The honey-yellow phosphate shows in the upper margin a tent-like fabric, indicating that the fast phosphatisation may have permineralized the soft tissue of the sponge.

**Fig. 7.**
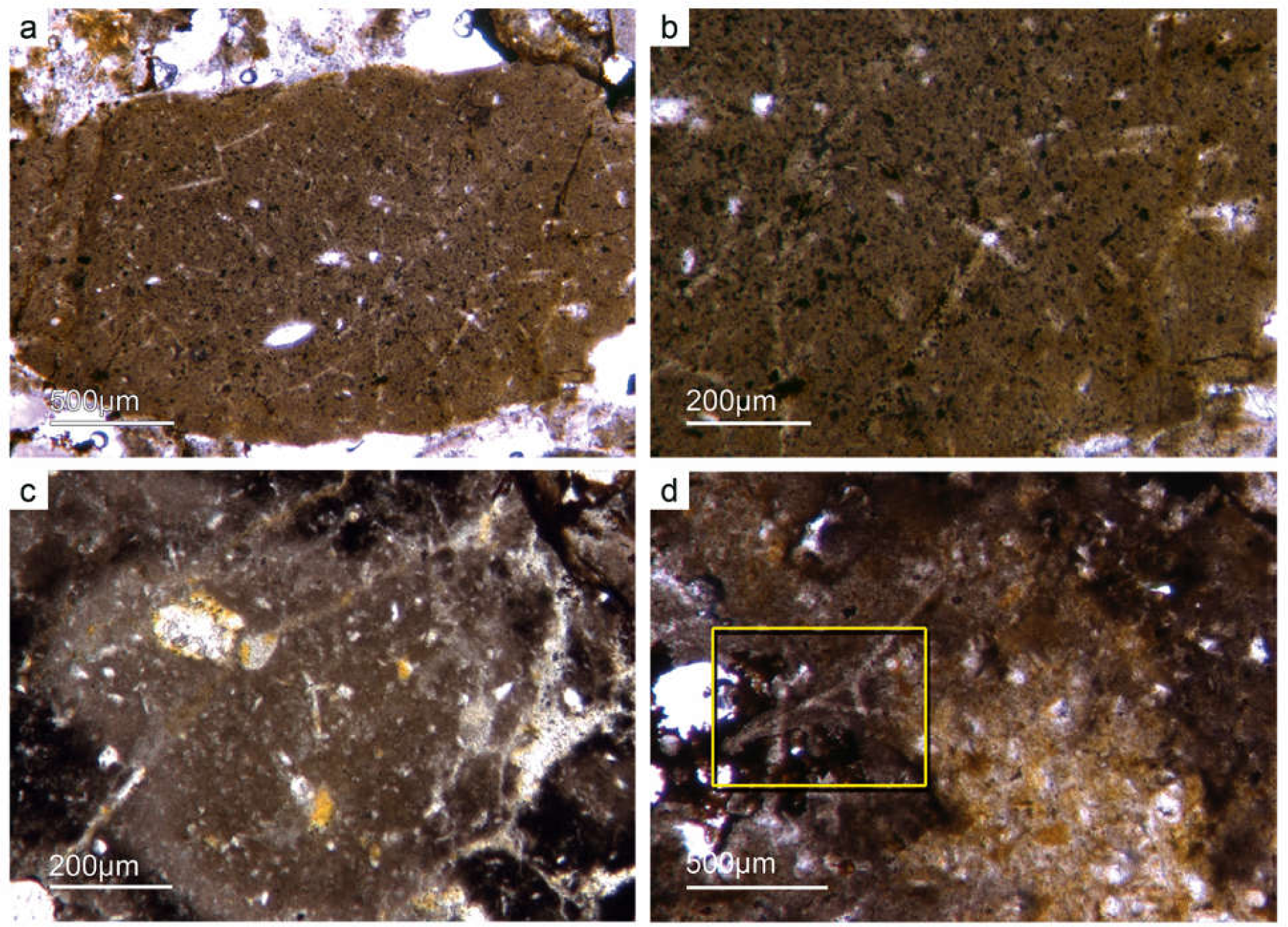
Lyssacine hexactinellid skeletons. (a) Parenchymal hexactinellid spicules within fine grained dark-brown phosphate (Fon27). The dermal areas of the sponge are lost. The phosphatic material shows abundant dark spots and grains probably former framboidal pyrite. (b) Detail of Fig. 7a demonstrating hexactin and monaxon parenchymal spicular skeleton. (c) Another lyssacine hexactinellid with spicules *in situ* (Fon20). The phosphate shows different grey colors, here interpreted as rapidly permineralized sponge tissue. (d) Remain of a more complex hexactinellid spicule, which is know from dictyonal skeletons (Fon98).

**Fig. 8.**
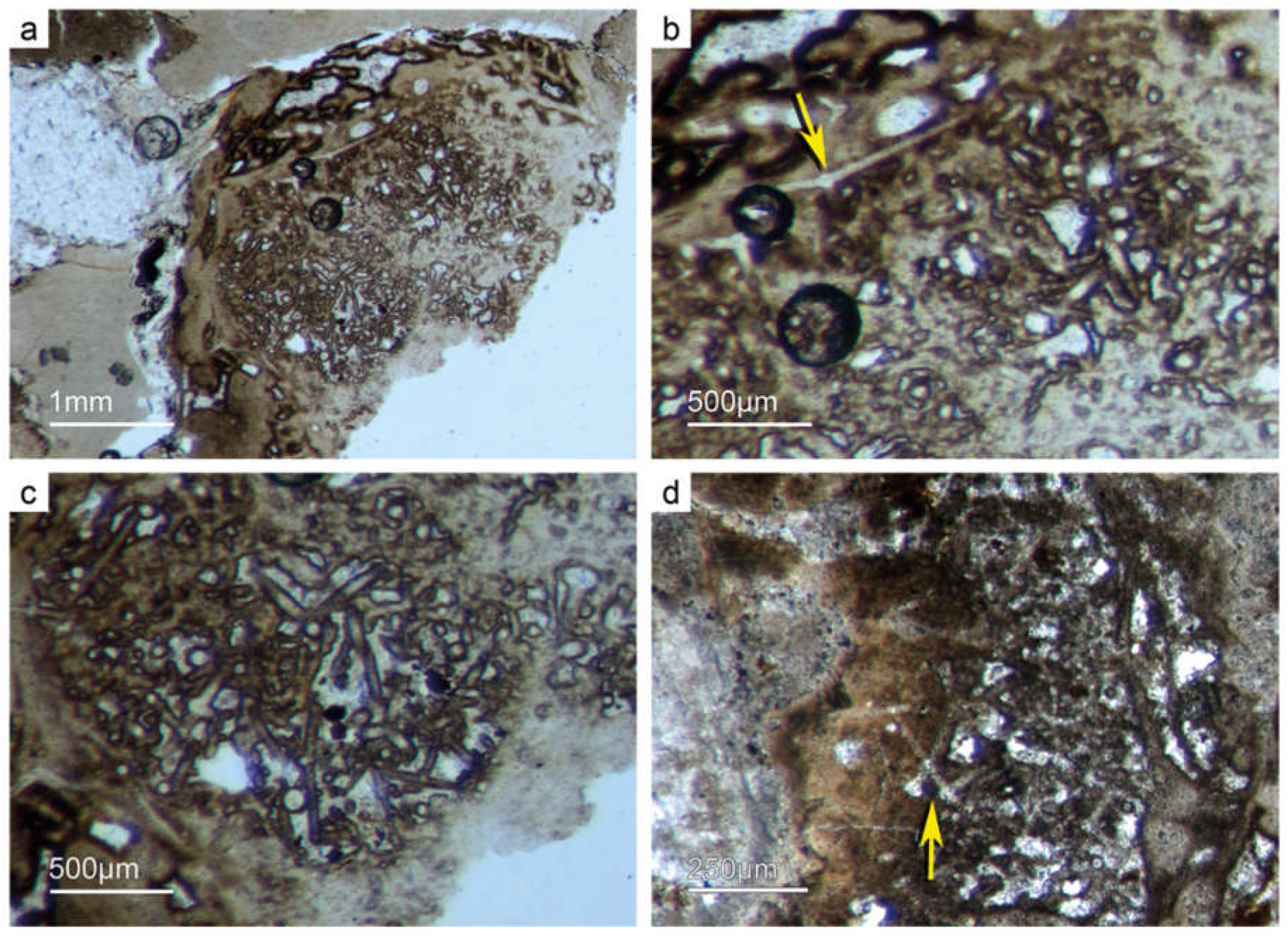
“Dictyonal” hexactinellid skeletons. (a–b) Nearly entirely preserved hexactinellid sponge with a cemented parenchymal and dermal spicular skeleton (Fon29). The marginal portions completely impregnated by honey-yellow phosphate and show ghost-remains of spicules and permineralized tissue remains. Preserved are large dermal spicules like stauractins (arrow in Fig. 8b). The spicules heavily cemented by phosphate and it is not clear if it is a real dictyonal skeleton – in any case a result of a very fast P-mineralization. (c) Another example of a parenchymal quasi dictyonal spicular skeleton formed by large hexactins surrounded by dark organic rich phosphate crusts. (d) This dictyonal skeleton exhibits a more primary spicular cementation and less phosphatic cements (Fon44).

**Fig. 9.**
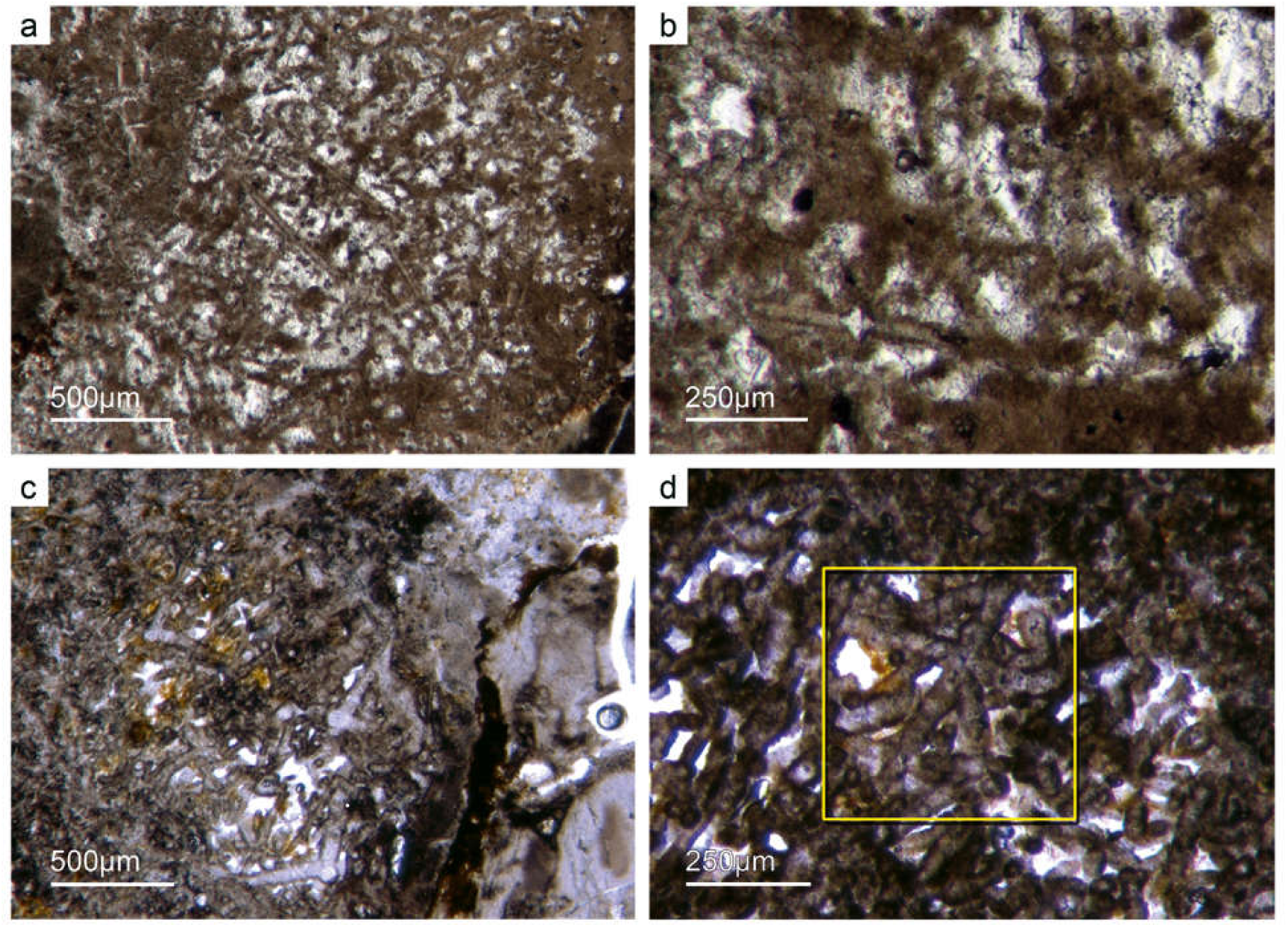
“Dictyonal” hexactinellid skeletons. (a–b) In some cases most of the spicules are replaced by brownish phosphatic material (Fon12). Space between is cemented by bright phosphatic cements. Some of the larger spicules also preserved in bright phosphatic cements but surrounded by dark phosphatic crusts as seen in Fig.8 the spicules fused together and forming articulated skeletons. (c–d) Best preserved “dictyonal” skeleton with large and small parchenchymal hexactin spicules and pentactin dermal spicules (Fig. 9d) (Fon26).

**Fig. 10.**
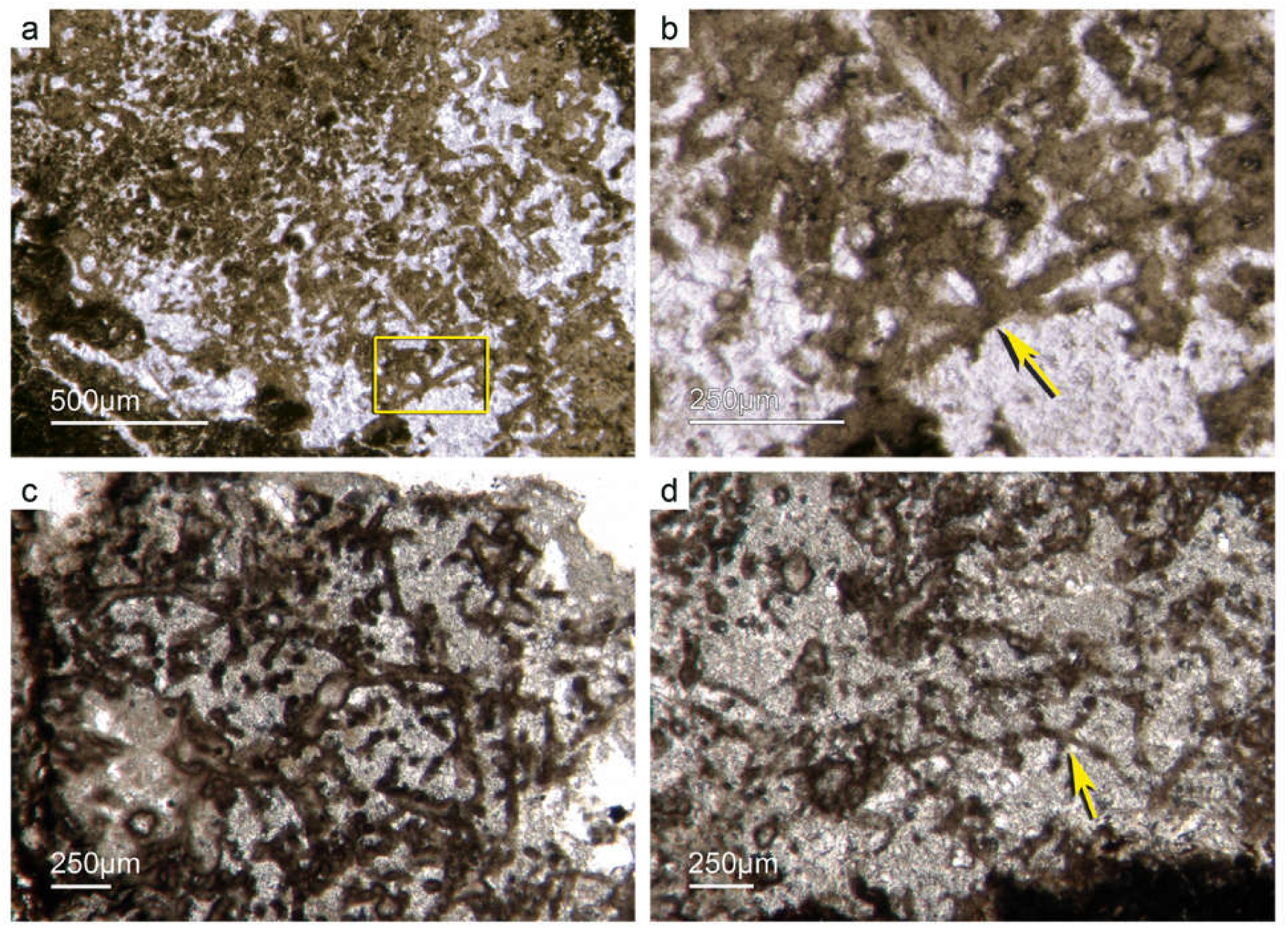
“Dictyonal” hexactinellid skeletons. (a–b) Primarily articulated fused spicular skeleton, mainly preserved in brownish phosphate (Fon16). Space between spicules cemented by bright phosphate. Remains of dermal skeleton exhibiting fused pentactins (Fig. 10b) (rectangle and arrow). (c–d) Dictyonal parenchymal hexactinellid skeleton (arrow) (Fon20).

The lyssacine morphotype of nearly articulated hexactinellids displays only weakly developed regularities in spicule arrangement. In few cases, only pentactins and stauractins are located at the sponges’ marginal dermal areas (“*Dermalia*” *sensu* Schulze, 1887). The basic hexactins (“*Principalia*” *sensu* Schulze, 1887) are randomly orientated and not attached within the lyssacine sponge tissue (Figs. 6, 7).

This is in contrast to almost entirely preserved hexactinellids that have fused spicule arrangements (Figs. 8–10). The choanosmal spicules (*Principalia*) are fused as in dictyonal skeletons. However, they do not exhibit a rectangular architecture known from other Dictyospongidae. It is possible to differentiate a few large (0.5 mm) fused hexactins among the abundant smaller ones (250µm). The dermal area exhibits large pentactins and tangentially arranged stauractins (*Dermalia*). These sponges are strongly impregnated with light brownish phosphate that likely replaces the primary sponge tissue. The spicules are often surrounded by dark, dense prismatic phosphate, and the inner parts are filled with a bright phosphatic cement (Fig. 8). A second type of preservation displays a brownish phosphatic spicule replacement, though a few have bright phosphatic cements. We observed a typical dictyonal spicule arrangement in a few choanosomal spicules. It is not clear whether this type of preservation is related to rapid phosphatization or it is a biologically mediated cementation (Figs. 8–9). If these spicular skeletons are pristine, it would be the oldest appearance of an advanced rigid hexactinellid skeleton.

### 5.c. Phylogenetic status of Demospongiae (Sollas, 1885)

Demosponges and hexactinellids are phylogenetically closely related and form a monophylum (e.g. Wörheide *et al*. 2012). The basic demosponge spicule types are monaxons and a four-rayed spicule called caltrops. The caltrop is the most characteristic demosponge stem group spicule (van Kempen, 1990; Reitner & Mehl, 1995, 1996; Reitner & Wörheide, 2002) (Fig. 11). Hexactinellids also harbor monaxon spicules. In living sponges, it’s possible to distinguish between hexactinellid and demosponge monaxons by checking the shape of the cross section of the axial filament: it is quadrate in hexactinellids and triagular or hexagonal in demosponges (e.g. Hartman, 1980; Reitner & Mehl, 1996; Leys *et al*. 2007). In the fossil record, this structure is difficult to be preserved pristinely. However, demosponges have a great diversity of monaxonic spicule types that differ from hexactinellid monaxons.

**Fig. 11.**
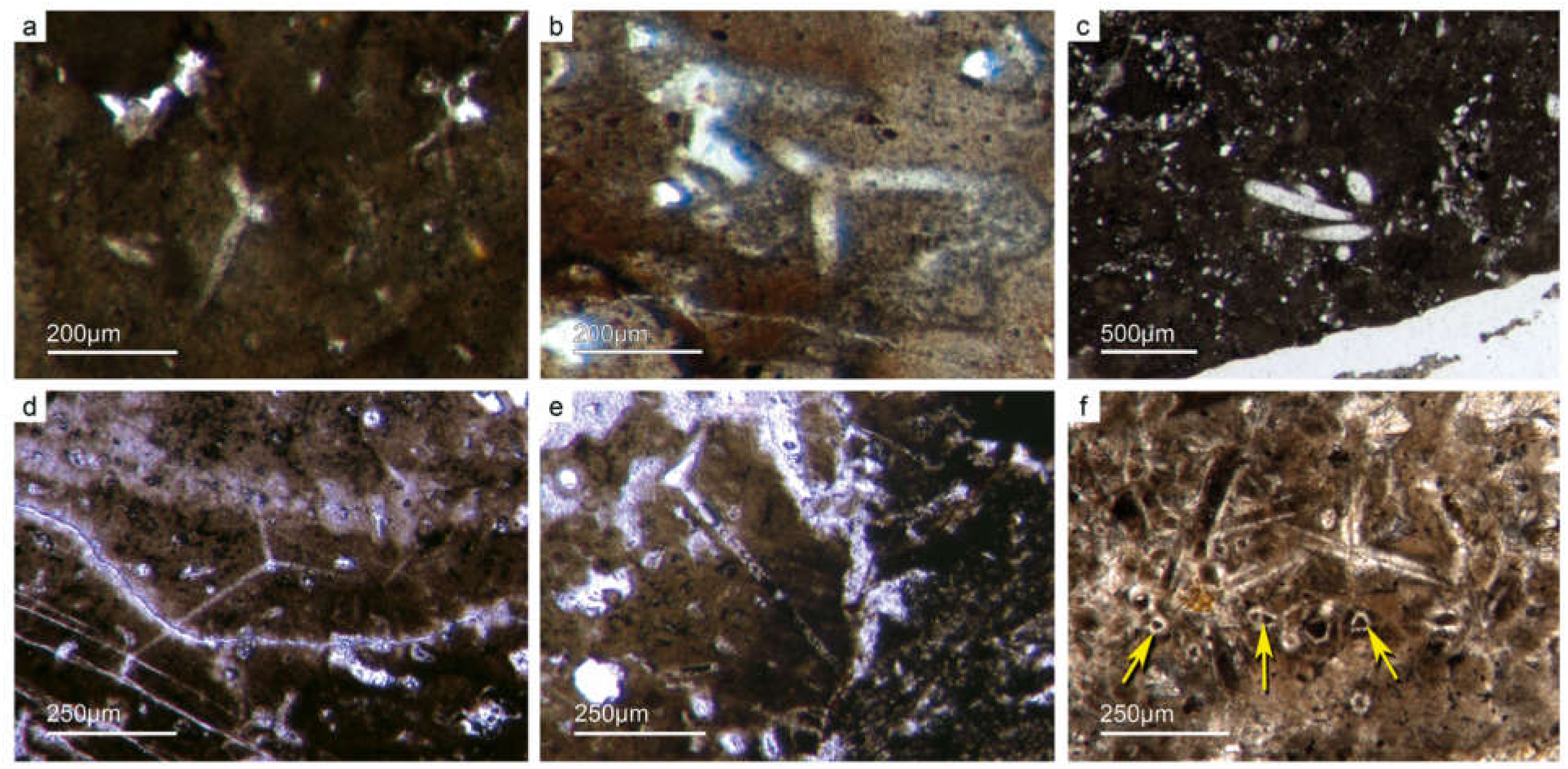
Basic demosponge spicules. (a–b, d) Various types of 4-rayed caltrop spicules with an angle of 120° between the rays (Fon31,59,11). (c) Group of thick monaxon style spicules (Fon11). (e) Typical dermal orthotriaen spicule (Fon11). (f) Aggregates of spicules with remains of the axial filament structure. Axial canal filled with organic matter (kerogen) and exhibiting hexagonal cross sections as known from demosponges (Fon10).

Furthermore, both sponge clades use roughly the same process to form spicules, with a proteinaceous fiber (axial filament) functioning as an organic matrix, mineralizing amorphous hydrated opaline silica. However, the fiber’s proteinaceous composition in demosponges is formed of silicatein α, a cathepsin L-like lysosomal proteolytic enzyme (Shimizu *et al*. 1998). In hexactinellids, cathepsin is observed more frequently than silicatein; in only one case a modified silicatein (AuSil-Hex protein in *Aulosaccus schulzei*) was detected (Veremeichik *et al*. 2011). In hexactinellids, collagen is also an important spicule seconary protein (Uriz *et al*. 2003; Müller *et al*. 2007; Ehrlich, 2010). This is in contrast to the sponge clade Calcarea which secrete calcitic spicules extracellular and not around an organic fiber (Jones, 1970). Spicule formation in sponges were developed phylogenetically independently multiple times.

Notably, various demosponges are non-spicular, and such forms were formerly described as “Keratosa,” or “horny sponges” (Minchin, 1900; Lévi, 1957). Recent phylogenomic studies have separated the non-spicular taxa into the clade “Keratosa” (Dendroceratida and Dictyocertida) and the clade Myxospongia (Chondrosida and Verongida) (Borchiellini *et al*. 2004; Erpenbeck *et al*. 2012; Wörheide *et al*. 2012; Redmond *et al*. 2013; Morrow & Cárdenas, 2015). The organic skeletons of these sponges are mainly composed of spongin, a composite of protein and/or chitin (Ehrlich *et al*. 2017). Spongin is still an enigmatic proteinaceous material that contains halogenated residues and until recently had not been sequenced (Ehrlich, 2019). This composite material is extremly resistent, as demonstrated by findings of chitin in middle Cambrian vauxiid keratose sponges (Ehrlich *et al*. 2013). Luo & Reitner, 2014 showed, for the first time, that “keratose” sponges are abundant in Phanerozoic fossil record, due to their special spongin-chitin skeletal taphonomy.

### 5.d. Fontanarejo spicule record of basic Demospongiae

#### 5.d.1. Non-articulated demosponge spicules

Different types of simple isolated caltrops were found within the Fontanarejo phosphorites (Figs. 11a, b, d). In a few cases, triaene dermal spicules, which are modified caltrops (Fig. 11e), were recorded. Large, thick spicule styles (ca. 0.5–1 mm long; 50–100 µm thick) were abundant and often arranged in bundles (Figs. 11c, 12a–d). Most of the spicules are embedded in yellow-brownish phosphorite, and often held together by a dark rim of phosphatic cement (Figs. 12a, d). Some of the spicules exhibit a clear hexa- to triangular central part formed by organic matter. This part could be the remains of a spicule’s central fiber, however, the preservation is unusual (Fig. 11f)

**Fig. 12.**
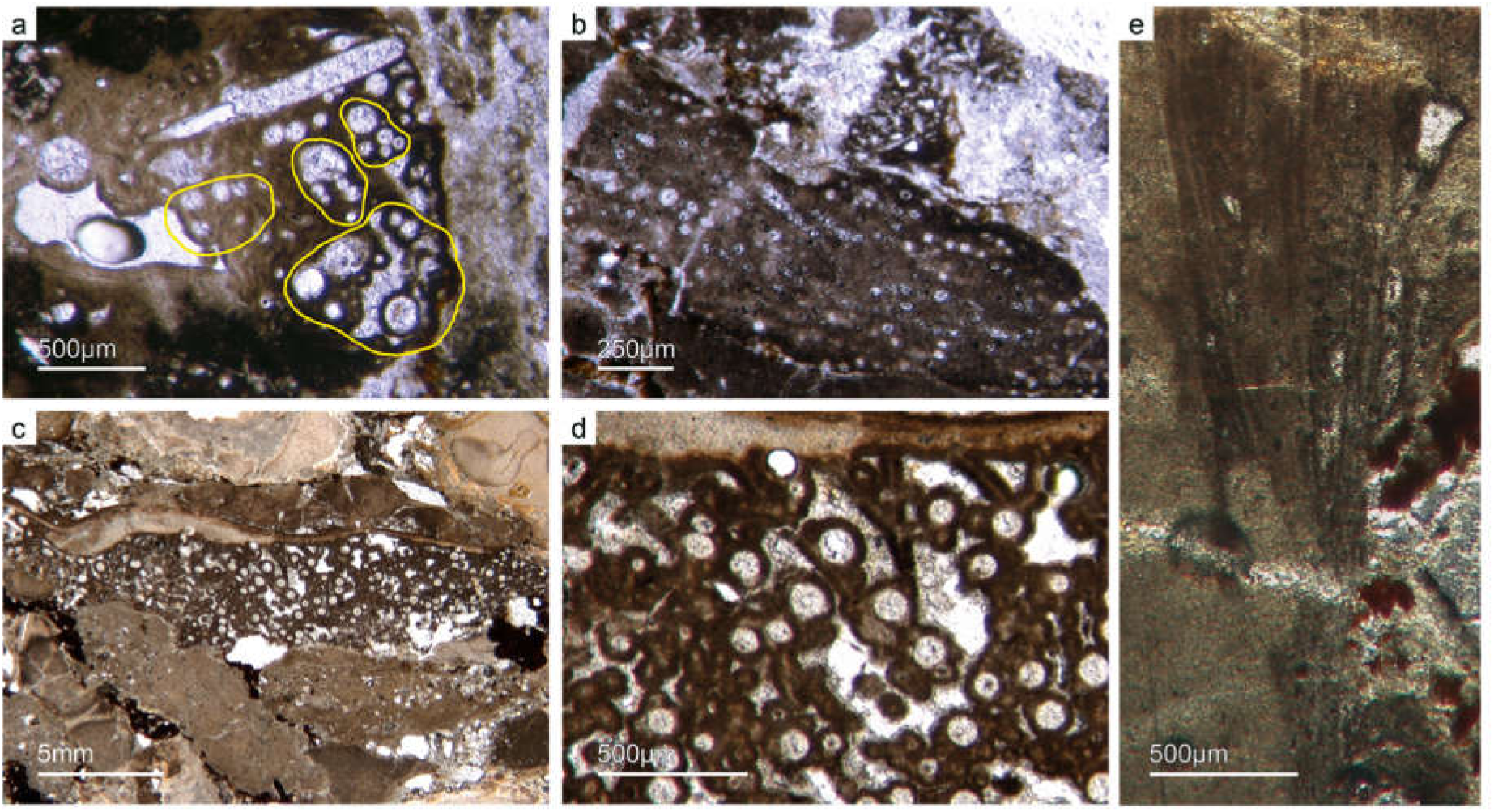
Articulated demosponge spicular skeletons. (a) Plumose bundle of large styles (Fon1). This arrangement of spicules known from the axinellid-halichondrid group. (b) Cross sections of styles mostly orientated in rows - probably part of dermal skeleton (Fon1). In (a–b) the sponges are preserved in honey-yellow to brownish phosphate, representing permineralized soft tissue. (c–d) Cross sections of monaxon spicules grouped in bundles (Fon4). The spicules surrounded by dark phosphatic cements, which keep them together. The arrangement of upright spicule bundles known from early Cambrian sponges like *Halichondrites* and *Hamptonia*. (e) A typical hamptonid spicule bundle formed by very long thin styles. General spicule architecture known from the axinellid-halichondrid group (Fon2019, 0007).

#### 5.d.2. Axinellid-halichondrid Spicule Architectures

The plumose arrangment of some of the spicule bundles resembles an ancient “axinellid” bauplan (e.g. Reitner, 1992; Reitner & Wörheide, 2002). In one sponge specimen, the spicule architecture is well preserved. The plumose spicule bundles represent outer dermal parts of a demosponge; the inner parts are constructed of tangentially orientated monaxons (Fig. 13e). Figs. 12e & 13d show another example of a plumose arrangement of spicules. The long, thin styles are nearly radially arranged, a structure known from lower Cambrian taxa such as *Choia*, *Choiaella*, *Hamptoniella*, *Hamptonia*, *Halichondrites*, and *Pirania* (Reitner & Wörheide, 2002; Finks & Rigby, 2004; Rigby & Collins, 2004; Botting *et al*. 2013). These taxa are all related to an axinellid-halichondrid bauplan. In some cases, bouquet-like arrangements of long styles (some millimeters) are observed close to a plumose arrangement of spicules (Fig. 13d). These bouquets types are present in ball-shaped modern taxa of Tethyidae and closely related with Hemiastrellidae (Hooper, 2002; Sará, 2002; Sorokin et al 2019). A very intriguing, almost entirely preserved, demosponge exhibits a well-developed, very dense etcosomal spicule architecture composed of short strongyles (c. 50 µm) (Fig. 14). The strongyles are interwoven and display a prefered tangential orientation (Figs. 14b, e). The endosomal spicules are large, thick styles (c. 200–300 µm long, 50 µm diameter) in plumose arrangement surrounded by long (200–400 µm), thinner styles (Figs. 14a, c, d). Some accessory curved monaxons are also present. This morphological character is relatively similar to an axinellid-halichondrid sponge type, but the ectosomal dermal layer is unusual.

**Fig. 13.**
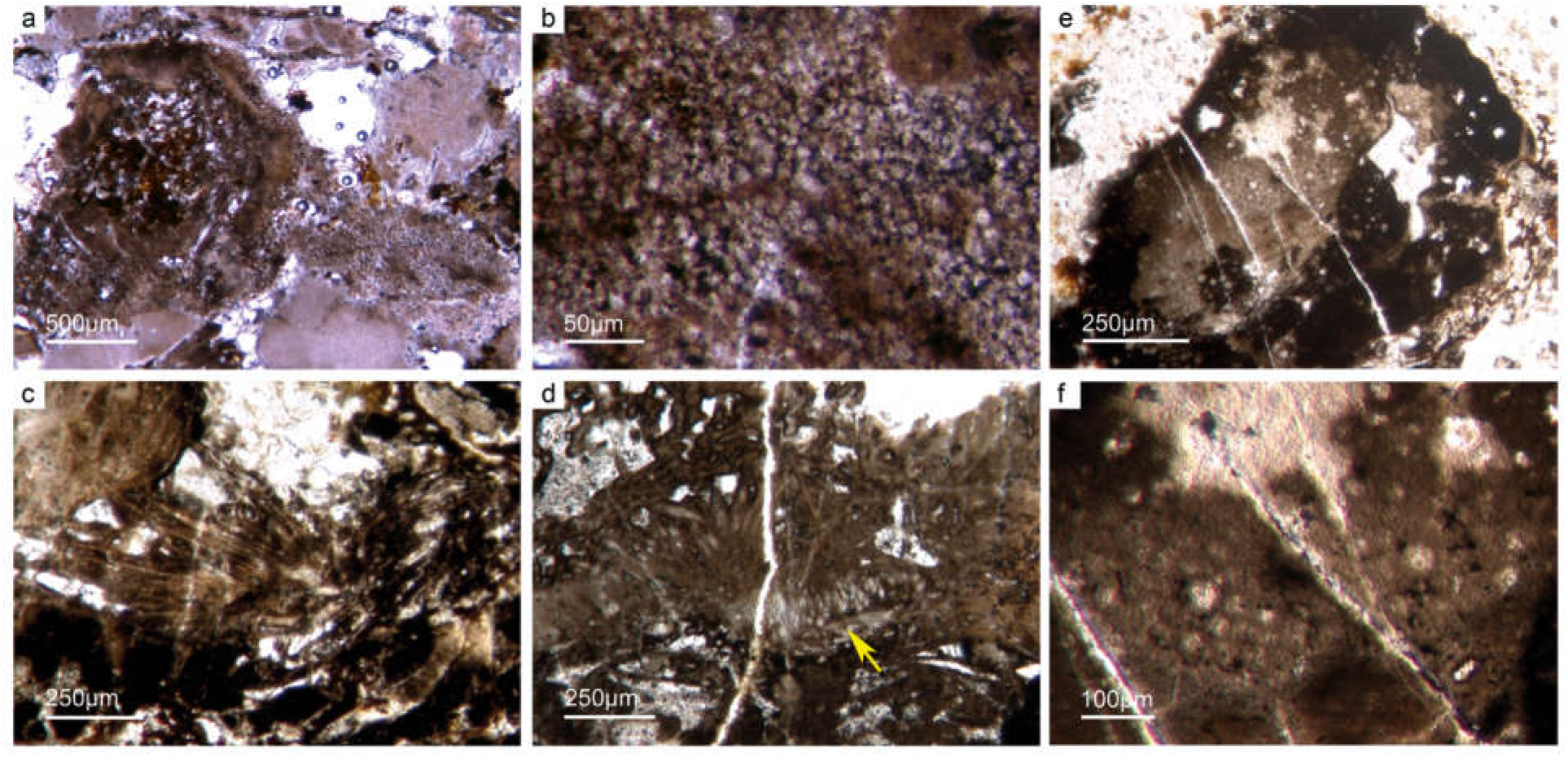
Articulated demosponge spicular skeletons. (a–b) Representing a “hadromerid” skeleton. The sponge shows a dense crust of putative asterose dermal microsclers and randomly orientated choanosomal, probably monaxon spicules (Fon 38). (c–f) The sponge represents again dermal asterose microsclers characteristic for chondrillid hadromerids (Fon17). (d) Radially orientated long styles typical for the early Cambrian sponges like *Choia, Choiaella, Hamptoniella* (Fon20). (e) Plumose arranged dermal short styles laying on inner tangentially arranged monaxons (arrow) (Fon20).

**Fig. 14.**
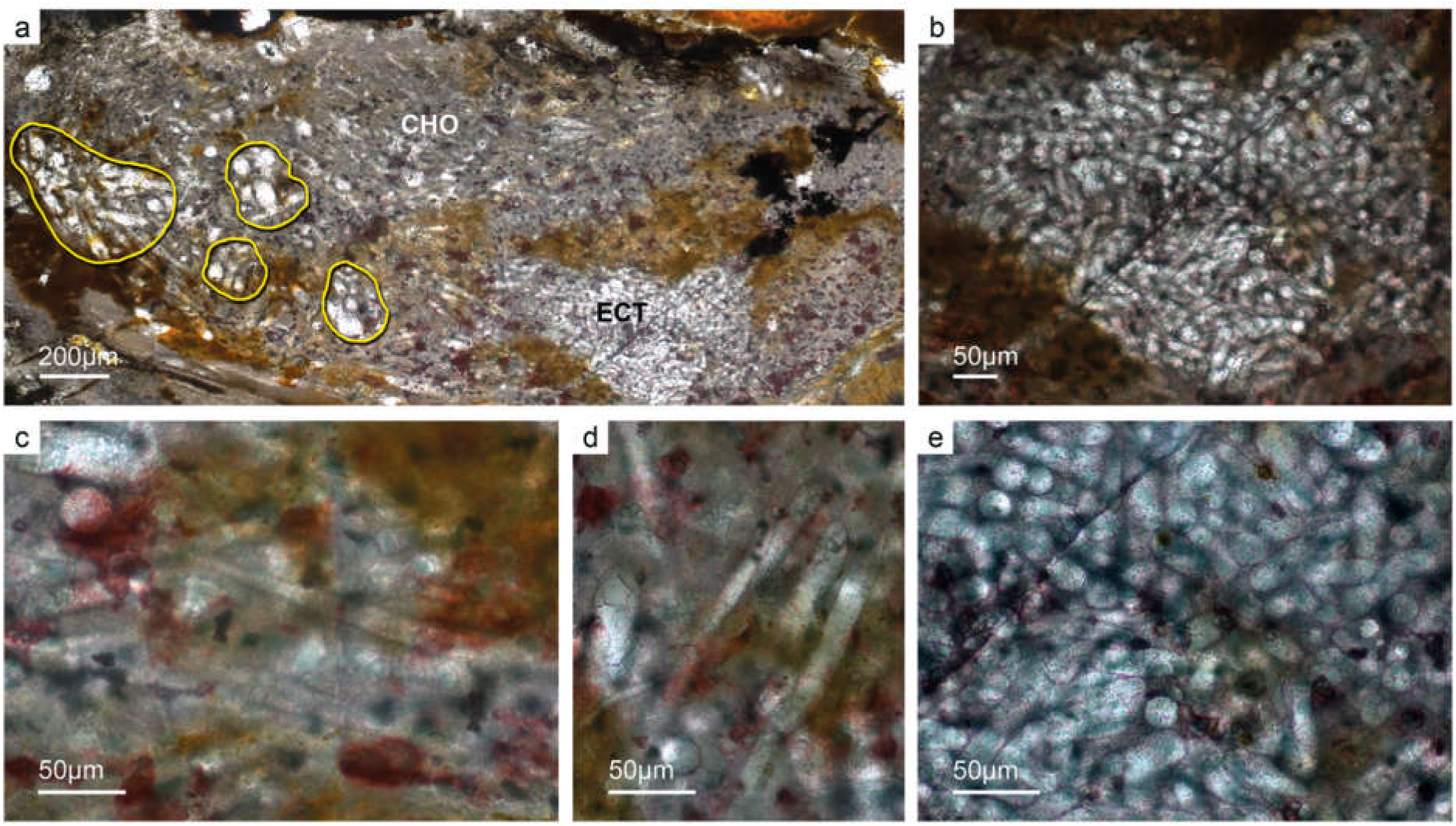
Almost entirely preserved axinellid-halichondrid sponge exhibiting a dense dermal layer. It exhibits plumose arranged thick large styles (a) and smaller styles (c–d) supplemented with a dermal layer of interwoven short strongyles (b, e) (Fon41).

#### 5.d.3. “Hadromerid” spicule arrangements

In addition to these plumose spicule arrangements, demosponges were observed with a thick dermal layer of small ball-shaped microscleres (c. 10–15 µm), probably asterose types (Fig. 13a, b). The choanosomal part of these sponges exhibit randomly ornientated, poorly preserved monaxone spicules, probably styles. This bauplan is known from “tetractinellid”-, hadromerid-, and hemiastrellid demosponges. A hadromerid affinity is most probable. In one case, only a dense dermal crust of putative asterose microscleres is observed (Fig. 13c, f). This sponge resembles the modern hadromerid taxon Chondrillidae, including the Cretaceous fossil taxon *Calcichondrilla crustans* Reitner 1991, which only bears dermal asterose microsclers (e.g. Boury-Esnault 2002; Reitner, 1991; Fromont *et al*. 2008). The lower Cambrian probable chondrillid demosponge exhibits some conchordance with the Cretaceous taxon *Calcichondrilla* (Reitner, 1991).

#### 5.d.4. Putative “Keratose” demosponges

Non-spicular demosponges are a sister group of spicular demosponges (e.g. Maldonado, 2009; Erpenbeck *et al*. 2012; Morrow & Cárdenas, 2015). Recent investigations demonstrated a widespread fossil record exists due to the special taphonomic behavior of skeletal spongin and chitinous fibers (e.g. Luo & Reitner, 2014, 2016; Friesenbichler *et al*. 2018). The best-known fossil taxon is the middle Cambrian Vauxida from the Burgess Shale, also recently discovered in the lower Cambrian Chengjiang Biota (Series 2, Stage 3) (Luo *et al*. 2020). Chitin was detected within this genus and is in accordance with the skeletal composition of the Myxospongiae (Verongida) (Ehrlich *et al*. 2013). Within the Fontanarejo phosphorites, few sponge remains have a keratose affinity. The putative sponge skeleton exhibits an irregular network of fibers similar to those already known from younger Phanerozoic keratose fiber networks (Figs. 15b, d, 16). Most of the fiber networks are embedded in dark brown phosphate matrices. They often form connecting points, and in some cases the fibers are curved. The fibers are preserved as a bright phosphatic cement. A second type exhibits a less dense fiber network (Fig. 15d). In another case, a reticulate fiber network is preserved in brownish phosphate (Fig. 15b). Micro-peloids, a taphonomic end product of sponge tissue alteration (Reitner *et al*. 1995; Reitner & Neuweiler, 1995, Reitner & Schumann-Kindel 1997), fill the spaces between the fibers (Figs. 15a, c).

**Fig. 15.**
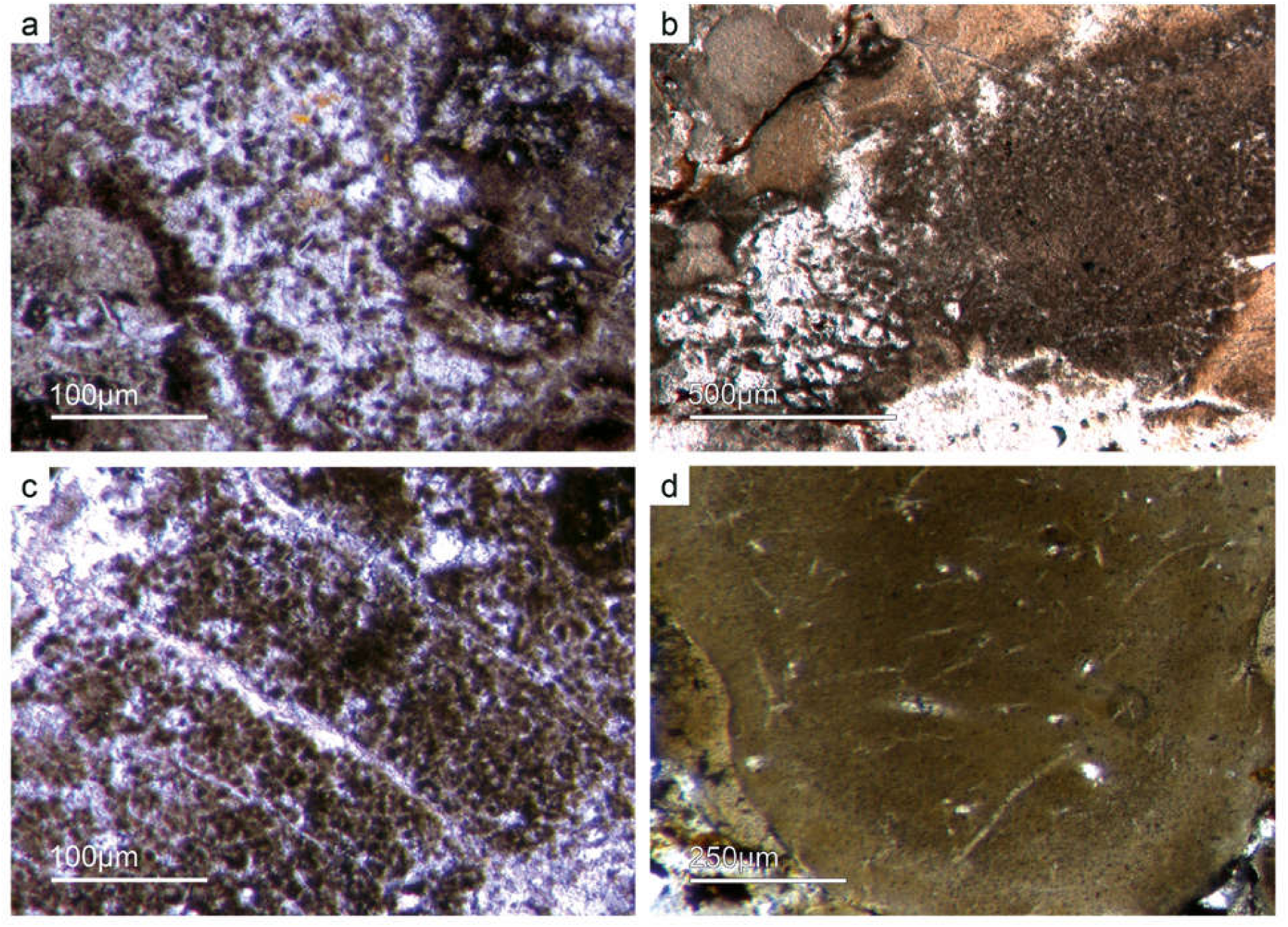
(a–c) Dark P-organic-rich micropeloids, a taphonomic end-product of microbial sponge tissue alteration (Fon16,13). (b) Non-spicular sponge skeleton with micropeloids between the sponge skeleton, interpreted as a “keratose” sponge affinity. (d) Within a brownish phosphatic clast thin filamentous fabrics are seen, interpreted as remains of sponging fibers (Fon56).

**Fig. 16.**
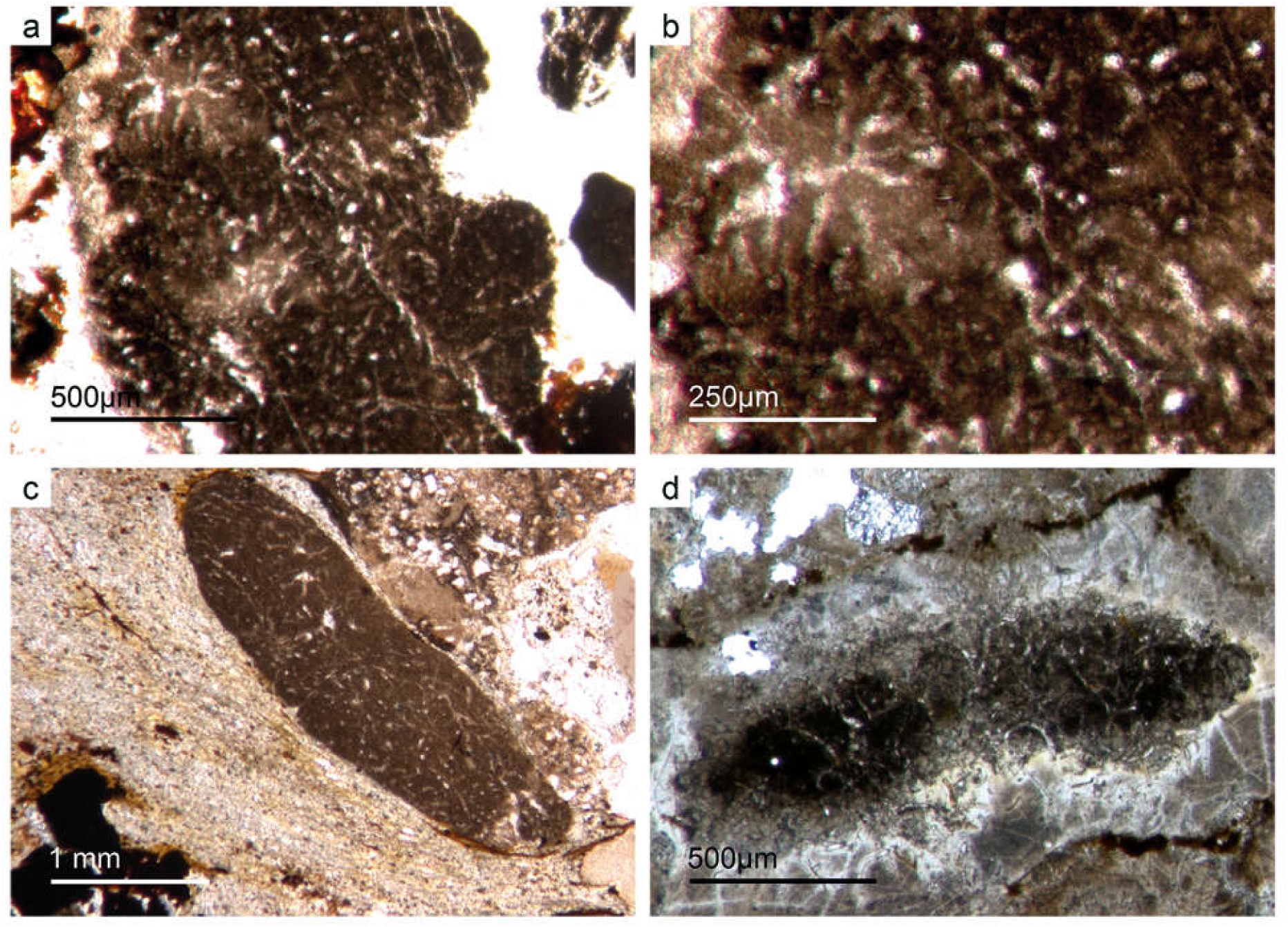
“Keratose” non-spicular demosponges. (a–b) Network of thin filamentous fabric within a dark-brownish phosphorite resembles a typical “keratose” organic skeleton (Fon77). (c) A phosphorite component with a clear fiber network (Fon19). (d) Dark phosphorite component with curved filaments partly forming a network (Fon76).

#### 5.d.5. Putative sponge larvae or resting bodies

Bundles of long styles with egg-shaped stuctures are observed within a few sponge “balls” (Fig. 17). These egg-shape structures have a diameter of c. 100–200 µm and resemble viviparous sponge larvae or gemmules. They have a distinct c. 10 µm thick bright rim that consists of radially orientated phosphatic crystals, a dark inner granular matrix, and small styles and/or diactine spicule remains. The dark inner granular matter is rich in iron hydroxides—probable remains of pyrite, and dark gray, irregular areas of phosphatic organic matter. The observed spicules are c. 20–50 µm long.

**Fig. 17.**
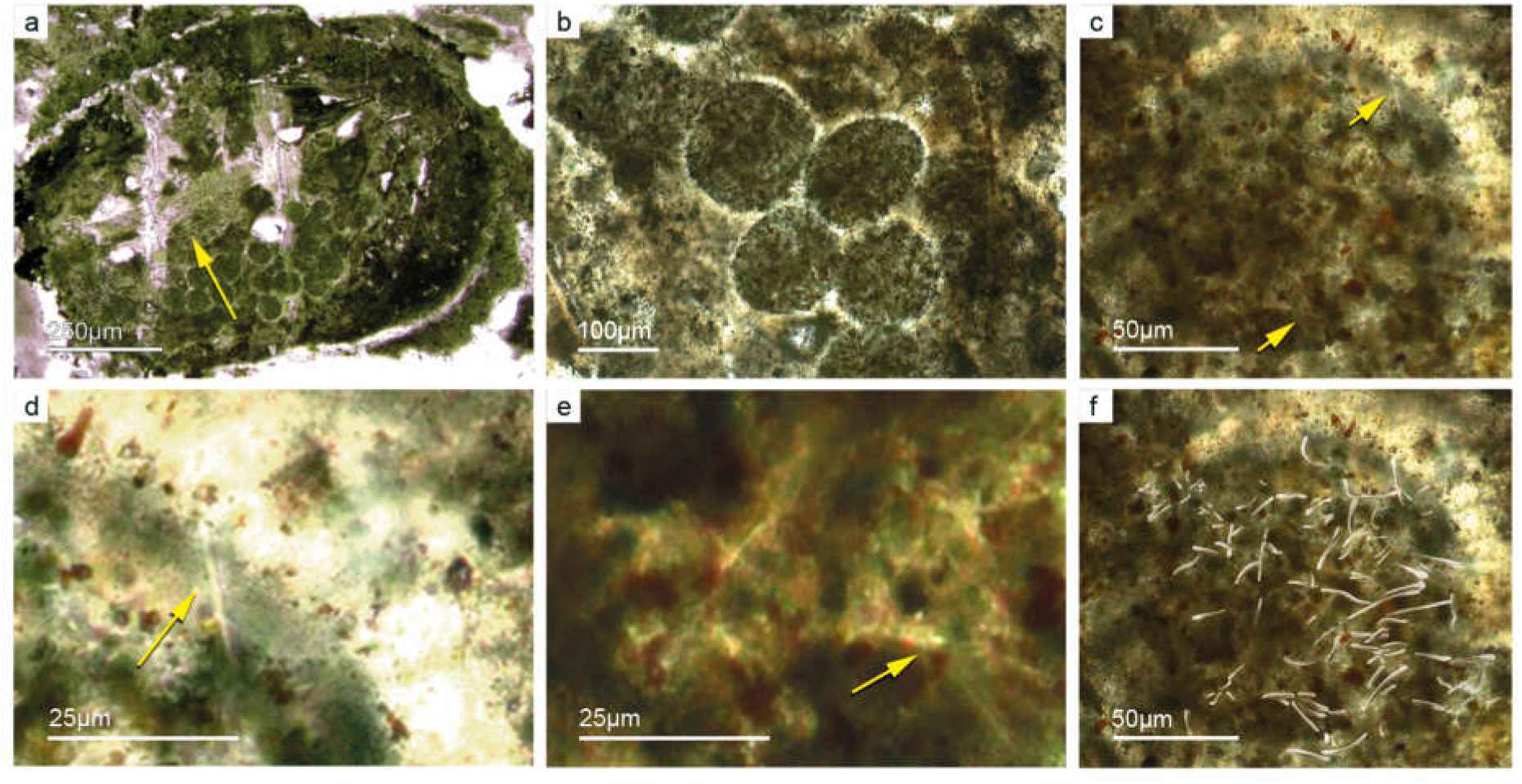
(a–b) Demosponge with a central bundle of styles associated with abundant egg-shaped structures (Fon 27). (c–e) These egg-shaped bodies show a distinct margin a central area with abundant Fe-rich small grains. Between the granular fabric are abundant small spicules (styles, oxea) (yellow arrows). In (f) recognized spicules are drawn. The egg-shaped bodies interpreted as larvae or resting cysts. These bodies show best analogy with demosponge parenchymella larvae.

The best analogue is the parenchymella of viviparous demosponges (e.g. from haplosclerids, poecilosclerids, and “Keratosa”), the only larva type with simple monaxonic embryonic spicules (Leys, 2003; Maldonado, 2006; Ereskovsky, 2010). To the best of our knowledge, fossil vivaparous sponge larvae (e.g. parenchymella-type) have not been observed until recently.

An alternative interpretation of the observed structures could be resting bodies like those of gemmulae. Gemmulae structures are very rare within the fossil record (Volkmer-Ribeiro & Reitner, 1991; Pronzato *et al*. 2017). Today, marine sponges with gemmulae are also rare and are restricted to a few taxa of the Haplosclerida (*Acervochalina loosanoffi*, family Chalinidae) and Hadromerida (*Protosuberites* sp.) (Connes *et al*. 1978; Arp *et al*. 2004). They are very common in freshwater sponges.

### 5.e. Putative small shelly fauna

As already discussed, determining the age of the Fontanarejo phosphorites is a central problem. As yet, no stratigraphically important fossils were found within the phosphorite-containing layers, thus the existing age was determined based on sedimentological correlation with nearby geological structures (Pieren Pidal, 2000; Jensen *et al*. 2010; Álvaro *et al*. 2019) and one zircon age published by Talavera *et al*. (2012). From the Alucida anticline, the early mollusk *Anabarella* was described, providing an important stratigraphic age (Pieren & García-Hidalgo, 1999; Vidal *et al*. 1999; Pieren Pidal, 2000). For the first time, we also found SSF remains in the Fontanarejo phosphorites, however, only in thin sections. The high amounts of siliceous cement make it impossible to etch out the microfossils. The various specimen slices observed in the thin sections are comparable to *Anabarella* (Fig. 18, Fig. 20a), and some tube fossils were also detected (Figs. 19a–b). These new SSF specimens provide convincing evidence that they are the same stratigraphic age as comparable specimens found in the Alucida anticline near Ciudad Real. One SSF exhibits a chambered structure comparable to phragmocones of cephalopods (Fig. 19c). The only known SSF with a phragmocone-like structure is the upper Cambrian monoplacophoran *Knightoconus* from Antarctica (Yochelson *et al*. 1973). Another option would be a chambered tube of “*Hyolithes kingi*” as described from the Lower Cambrian of Jordan by (Bandel, 1986). Mollusk-type SSF tube fossils are also present. In thin section we observe simple narrow shafts of some millimeters in length and 50–100 µm in diameter (Fig. 19a–b). Commonly, we observe cross sections of putative tube fossils with a diameter of maximum few millimeters in length, often exhibiting a week outer ornamentation. The shell structure is often prismatic (Figs. 20b–c).

**Fig. 18.**
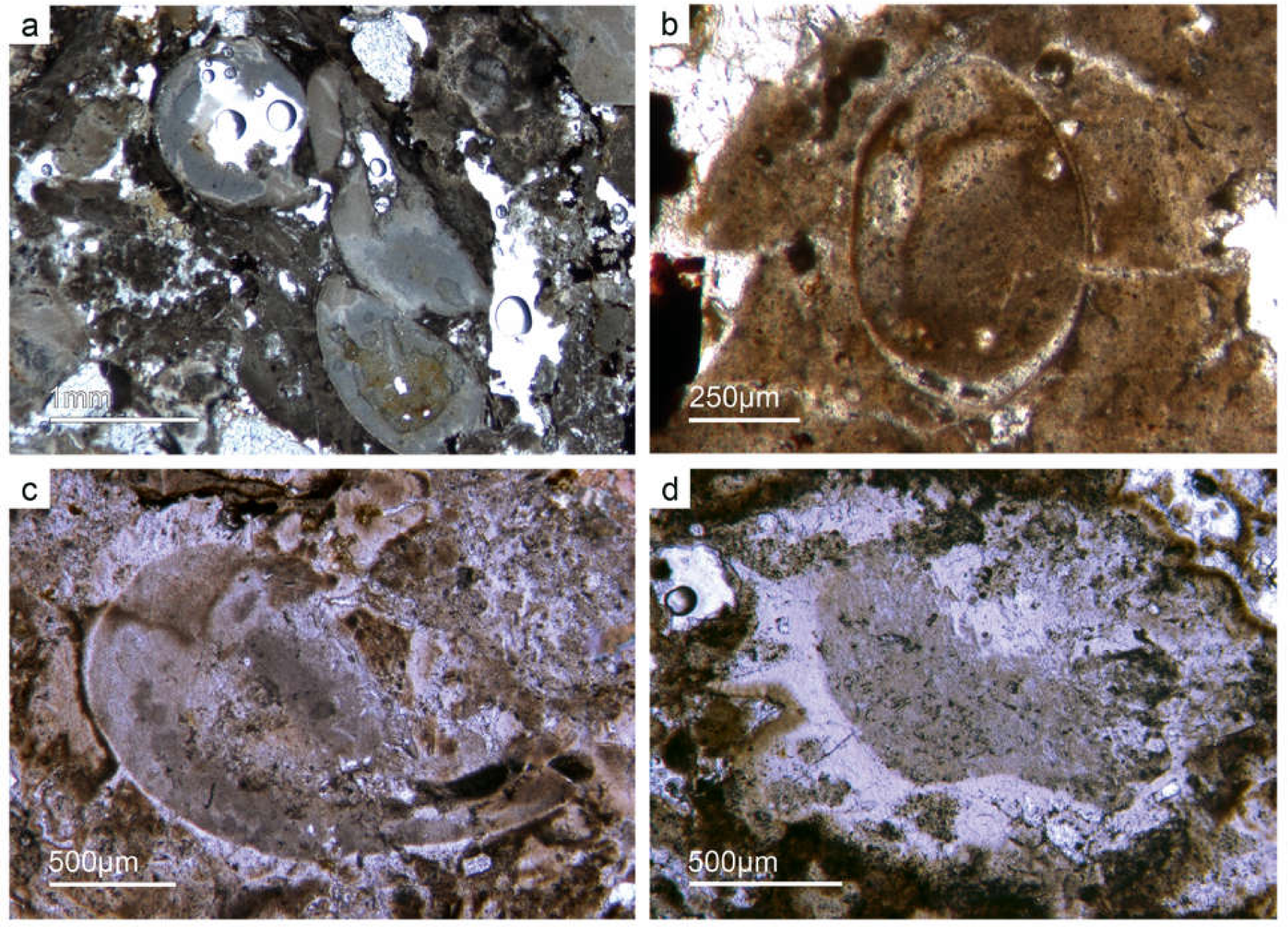
Mollusk remains (SSF). (a–c) Axial and oblique cuts of *Anabarella* sp., a putative monoplacophoran mollusk (Fon61,35). (b) Vertical cut of *Anabarella* sp. (Fon4), (d) Oblique cut of an ornamented *Anabarella* type (Fon35).

**Fig. 19.**
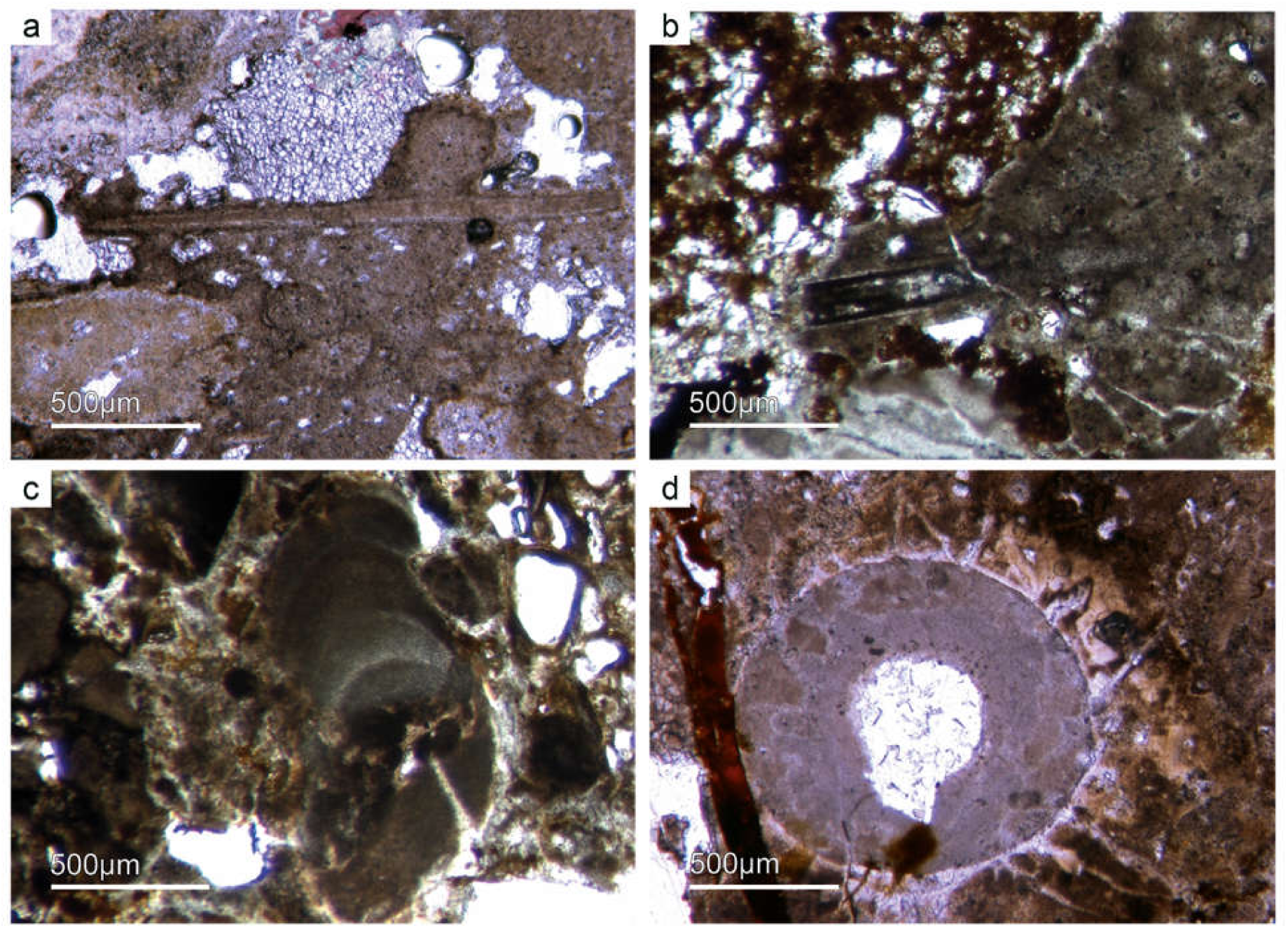
Multiple types of SSF. (a–b) A long thin tube shell (Fon99). (b) Another type of long thin tube shell (Fon44). (c) A new form of a chambered SSF (Fon31). (d) A round cross section of a tube fossil (Fon78).

**Fig. 20.**
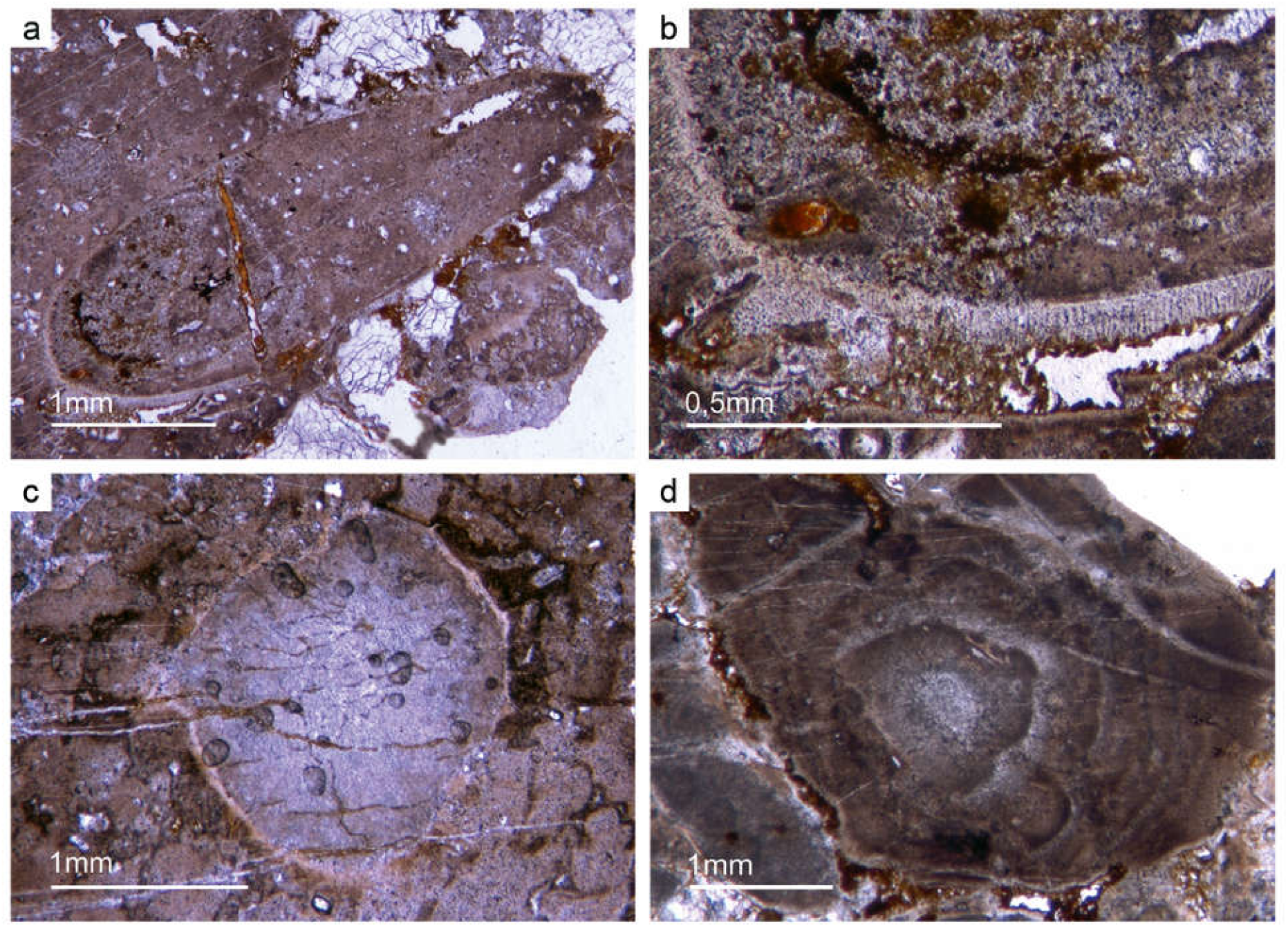
(a) A median-oblique cut of *Anabarella* sp. (b) A vertical cut an ornamented SSF. (c) Detail of the prismatic phosphatic shell structure of Fig. 20b (Fon4). (d) Median-oblique cut of a cloudy-colloform microbialite (Fon92).

### 5.f. Implications of the Fontanarejo record on early metazoan evolution

Molecular clock analyses suggest that sponges evolved in the Neoproterozoic. However, no convincing morphological fossils or spicules have been reported from that era. For this reason, the molecular clocks are usually calibrated with sedimentary hydrocarbons in Cryogenian and Ediacaran rocks and oils that are widely considered as being indicative for sponges (i.e., 24-isopropylcholestanes) (Sperling *et al*. 2010; Erwin *et al*. 2011; Cunningham *et al*. 2016; Gold *et al*. 2016; Dohrmann & Wörheide, 2017). This is because demosponges are the only extant organisms known to biosynthesize the precursor molecules of these distinct compounds as major lipids (e.g. Hofheinz & Oesterhelt, 1979; Djerassi & Silva, 1991; Giner, 1993; McCaffrey *et al*. 1994; Love *et al*. 2009; Zumberge *et al*. 2018). 24-isopropylcholestane precursors are also known as insignificant (that is, two orders of magnitude less abundant) by-products in the biosynthesis of 24-*n*-propylcholestane lipids by extant pelagophyte algae (Giner *et al*. 2009; Love & Summons, 2015). Minor traces of these compounds in modern rhizarian protists (Nettersheim *et al*. 2019) seem to be problematic and are subject of an ongoing discussion (Hallmann *et al*. 2020; Love *et al*. 2020).

Regardless of this debate, our finding of taxonomically diverse demosponge and hexactinellid communities in the late Fortunian Fontanarejo phosphorite further corroborates a Precambrian origin of the phylum Porifera. Notably, the organismic processes involved in biomineralization are enourmous complex, and there are various extant groups of sponges that seem to belong to the non-spicular lineages and do not exhibit a distinct body plan. For all these reasons, it appears likely that the most ancestral representant of the sponge phylum (or “Urschwamm”) solely consisted of soft tissue, perhaps rather resembling a microbial mat (Luo & Reitner, 2016). This would be in good agreement with the idea that microbial mats transitionally evolved into sponges, as hypothesized earlier (Reitner, 1998). And, such an scenario would explain the seeming mismatch between paleontological findings, the sedimentary hydrocarbon record, and molecular clock data.

## 6. Results and discussion: sedimentary evidence for microbial mediated phosphate formation

The abundant cloudy textures observed in the Fontanarejo phosphorite strongly suggest a microbial origin of the deposit, as suspected earlier by Álvaro et al. (2016). The microbial impact on sediment formation is further evidenced by a variety of components such as mm-sized aggregates of phosphatic material that exhibit irregular to cauliflower-like shapes and often contain small allochthonous grains as for instance siliciclastic components and spicules. Notably, these components represent up to 20% of the entire phosphatic sediment. A further example are abundant oncoidal-like components with roughly laminated colloform and thrombolitic structures and a distinct nucleus (Figs. 20d, 21). Raman spectroscopy revealed the presence of pure and organic-rich apatite, organic matter, as well as of accessory minerals such as anatase and quartz in these components (Figs. 22, 23). The laminated to thrombolitic structures and the close spatial association of organic-inorganic phases indicate that apatite formation was probably mediated *via* microbial exopolymeric substances (EPS) and/or fueled by phosphor deriving from bacterial cell degradation, perhaps in biofilms (see below).

**Fig. 21.**
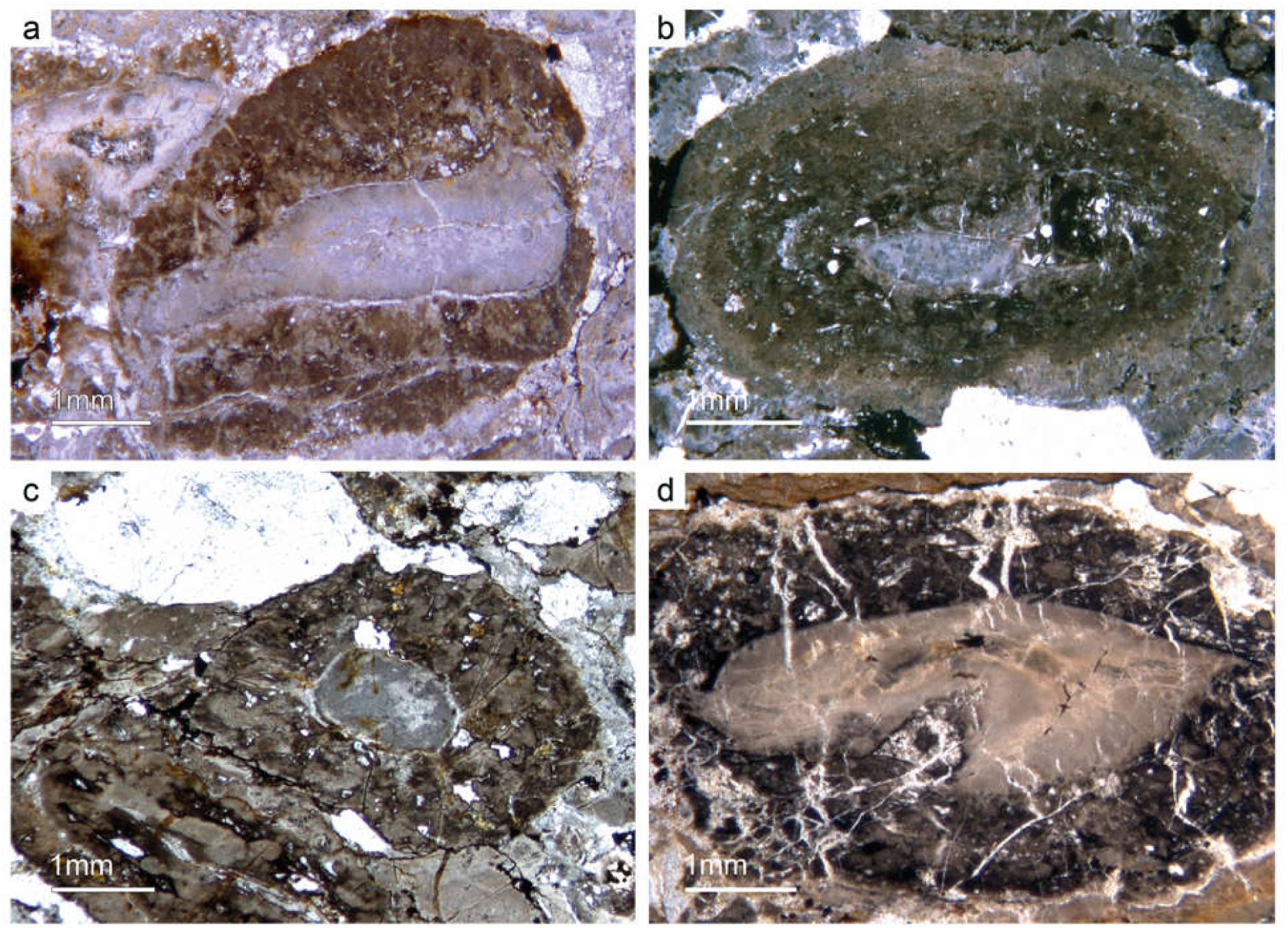
Oncoid-type microbialites. (a) Irregular cauliflower growth mode of phosphatic microbialite encrusting on a phosphatic clast (Fon18). (b) Roughly-thrombolitic growth structure overgrown a phosphatic core component (Fon35). (c) Thrombolitic oncoid-type microbialite (Fon44). (d) Dark organic-rich thrombolitic oncoid-type microbialite (Fon21).

**Fig. 22.**
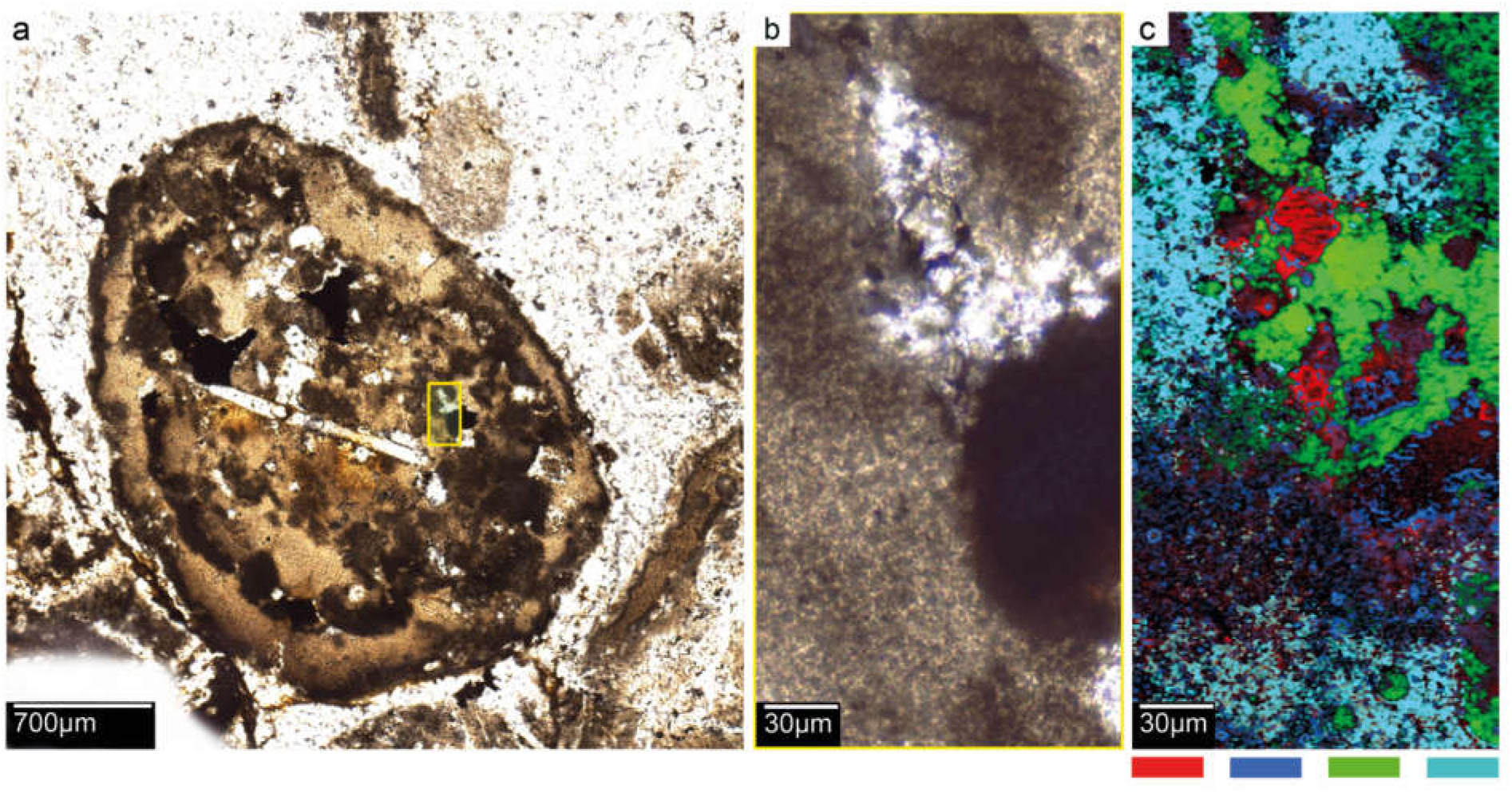
(a) Median-oblique cut of a cloudy-colloform microbialite, showing in the center a large stronglye megasclere. Rectangle marks the area of a Raman map. (b) Detail of the map area. (c) Raman map: red, organic matter; dark blue, TiO_2_ – anatase; green, apatite; blue, apatite + organic matter (Fon10).

A second type of microbialite are mm-sized micro-domal stromatolites. Generally, the structure of these stromatolites is similar to the colloform structure of the phosphatic oncoidal-like components described above. However, the stromatolites exhibit a distinct lamination of alternating, c. 50 µm thin dark-brownish, brownish and white layers (Fig. 23). The dark-brown and brown layers consist of organic-rich and organic-poor apatite, respectively. The white layers, in contrast, are composed of siliceous apatite or pure silicate. As in case of the oncoidal-like components (see above), the organic-rich apatite in the stromatolites is likely a product of EPS controlled mineral precipitation or taphonomic biofilm degradation. TiO_2_ is enriched in few layers and might be an authigenic, perhaps even microbially mediated precipitate; however, unfortunately these processes are still too poorly understood to draw robust conclusions (e.g.Taran *et al*. 2018; Izawa *et al*. 2019).

**Fig. 23.**
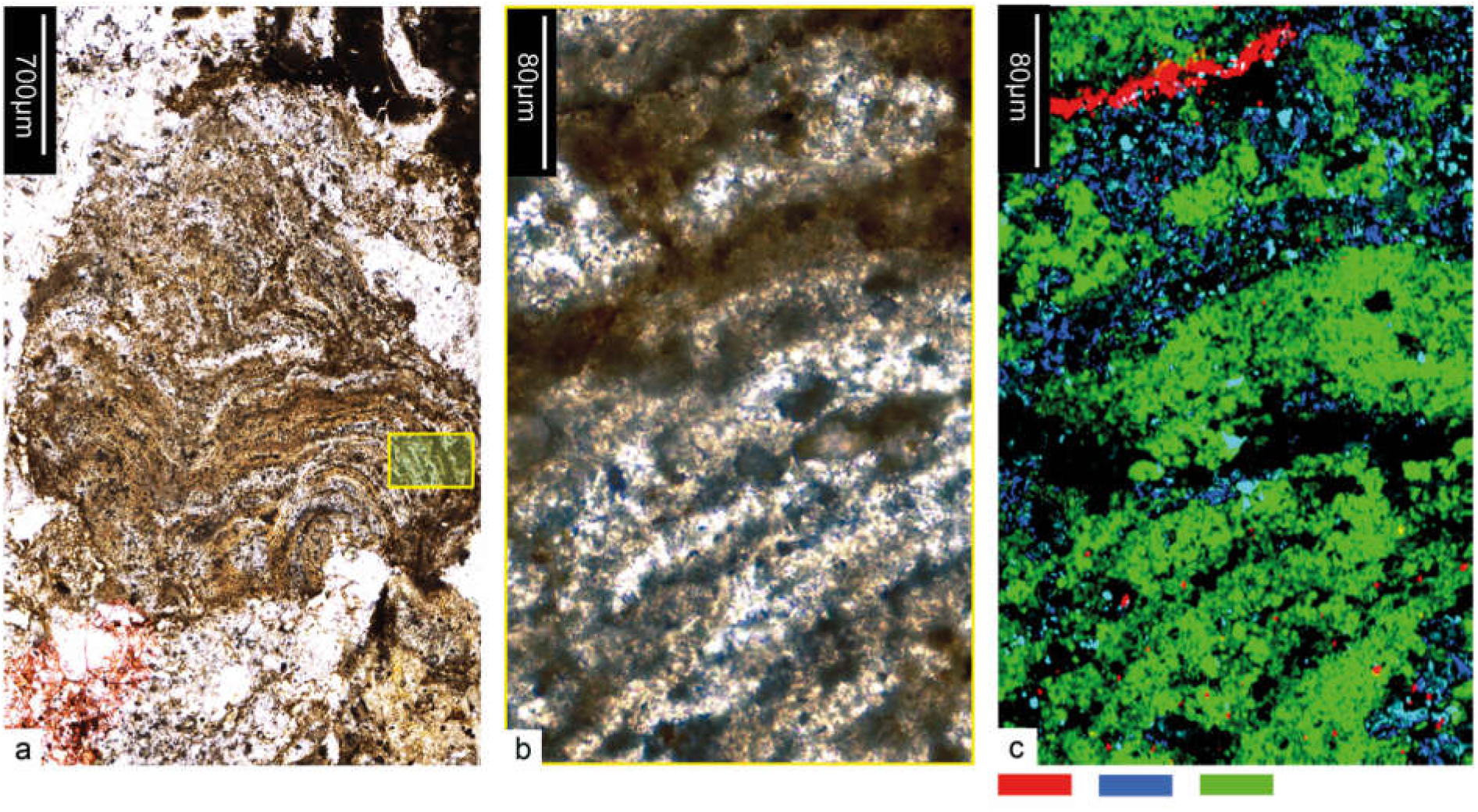
(a) A domal microstromatolite, rectangle marks the Raman map area. (b) Detail of the map area. (c) Raman map: red, TiO_2_ – anatase; blue, apatite + organic matter; green, apatite without organic matter (Fon10).

The presence of microbes in the Fontanarejo phosphorite environment is further evidenced by the presence of large filamentous fossils encapsulated in some of the observed microbialites (Fig. 24a-c). These fossils strongly resemble sheets of extant sulfide-oxidizing bacteria (SOX-B) (Peckmann *et al*. 2004; Bailey *et al*. 2013; Birgel *et al*. 2014; Andreetto *et al*. 2019). This interpretation has important implications for the formation of the Fontanarejo phosphorite deposit. Today, SOX-B are particularly widespread in sediments of coastal upwelling regimes off the west coasts off Africa and South America (Arning *et al*. 2009a,b). Notably, vacuolated SOX-B like the giant bacterium *Thiomargarita* (Schulz & Schulz, 2005; Brock, 2011) as well as filamentous SOX-B like *Beggiatoa* and *Thioploca* all harbor intracellular polyphosphate inclusions (e.g. (Jørgensen & Gallardo, 1999; Salman *et al*. 2011; Crosby & Bailey, 2012). It has been shown that the hydrolysis of the polyphosphates in *Thiomargarita*-rich sediments releases sufficient phosphor into the pore water to precipitate apatite (Schulz & Schulz, 2005). Moreover, it has been demonstrated by ^33^P-labeling experiments that phosphate taken up by SOX-B is rapidly transformed from intracellular polyphosphate to apatite (Goldhammer *et al*. 2010). All these studies support the hypothesis that microbial metabolism can foster apatite formation, and we consider this as plausible explanation for the formation of the Fontanarejo phosphorite.

**Fig. 24.**
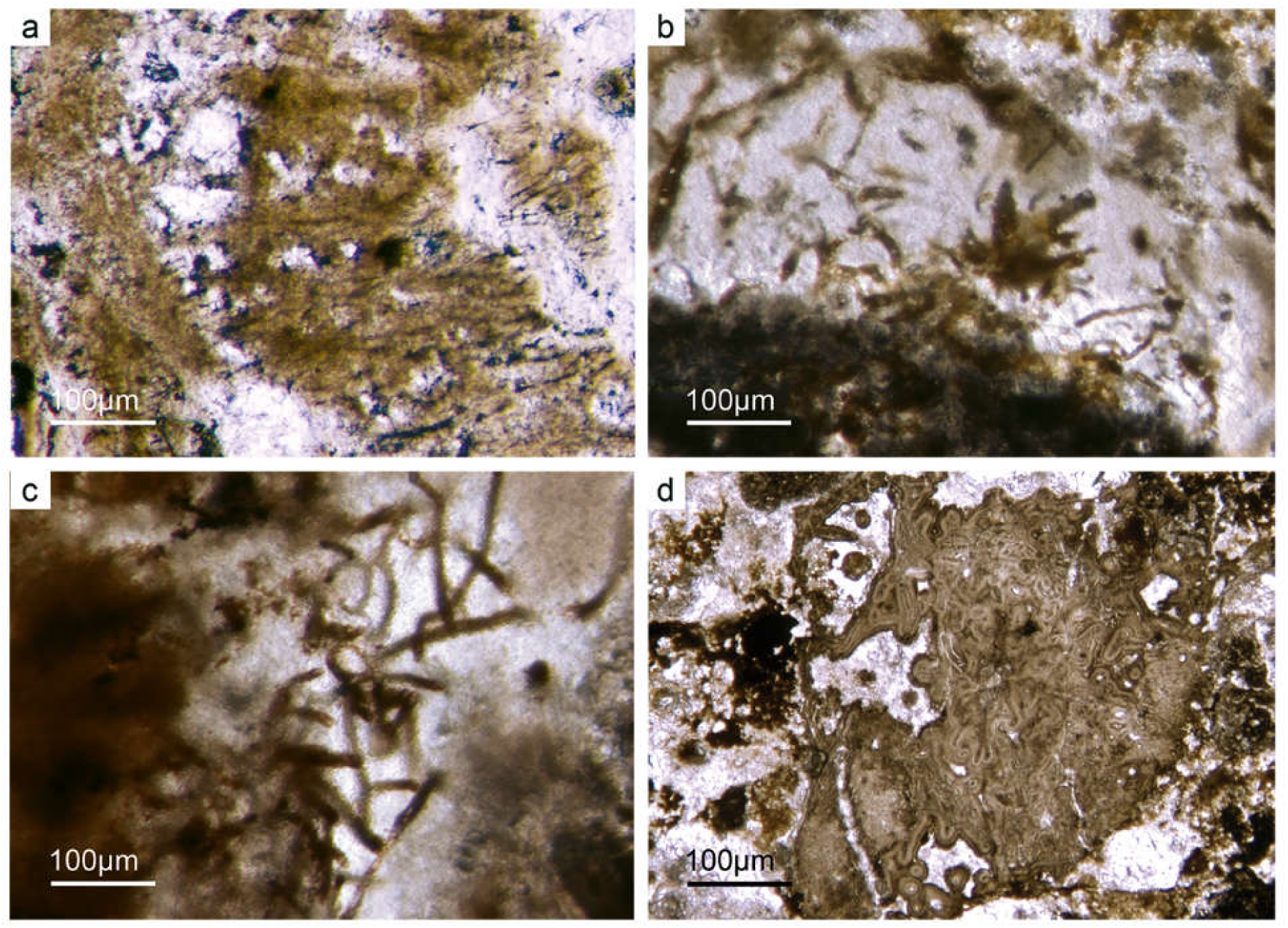
Filamentous microbial remains. (a) Long filamentous structures, probably remains of SOX-bacteria (Fon35). (b) Filamentous remains within bright phosphorite (Fon36). (c) Network of large filaments interpreted as remains of SOX-bacteria (Fon36). (d) Sponge skeleton heavily cemented by multilayered phosphate – a result of intense P-rich fluid flows.

In non-upwelling systems, apatite formation appears to be linked to a non-microbial process termed “iron-redox pump” (e.g. Einsele, 1936; Mortimer, 1941; Nürnberg & Peters, 1984; Ruttenberg & Berner, 1993). In anoxic pore water environments, iron(III) will readily be reduced to iron(II). As a consequence, iron(III) bearing mineral phases will dissolve, thereby releasing adsorbed phosphor into the pore water, which then further reacts to authigenic apatite. Intriguingly, the Pusa shales are locally black and enriched in pyrite, indicating low-oxygen conditions. These conditions would have been favorable for an “iron-redox pump”, which might account for some of the observed phosphatic cements (Fig. 24d). However, the abundance of evidence for microbial processes still strongly suggests a microbial origin for most of the observed apatite in the Fontanarejo deposit. In combination, both pathways plausibly explain the rapid precipitation of apatite (Cook & Shergold, 1984; Föllmi, 1996) which accounts for the exceptional preservation of microbial features and lyssacine and articulated sponge fossils.

## 7. Conclusions

1. The Fontanarejo phosphorites were described decades ago, but have not been studied in detail since then. A new detailed geological map is presented, updating and improving previous mappings. The stratigraphic and sedimentological aspects of these deposits have been controversial and are here reviewed and discussed, supporting the interpretation that they are late Fortunian in age and formed as channels of a turbidite system located at the platform-slope transition. Furthermore, we present the first microfacies description of thin sections from these deposits, and the first comprehensive characterization of their paleontological content, highlighting the relevance of their sponge fossils.
2. The phosphorites of Fontanarejo harbor abundant sponge remains. Hexactinellid remains are most abundant, including lyssacine and possible dictyonal spicular skeletons. In addition to basic hexactins, diactins, and dermal stauractins, pentactins are common. The spicules have simple morphology, and are not preserved with any microsclers. The lyssacine sponges are often preserved in honey-yellow phosphatic material that likely is permineralized soft tissue. They exhibit only a few parenchymal spicules and, in rare cases, stauractin dermal spicules. This type of hexactinellid is related to the “Rossellimorpha A” *sensu* Mehl-Janussen (1999), in contrast to the hexactinellids with dictyonal skeletons. The parenchymal areas show a very dense spiculation of fused hexactins. Hypodermal pentactins are present.
3. Demosponges are less common but show a higher morphological diversity. Isolated basic caltrop-spicules of the stem group demosponges, triactine dermal spicules, and diverse monaxonic spicules are common. Articulated plumose and bouquet-type spicule arrangements are related to the axinellid-halichondrid group. An unusual type of an axinellid-halichondrid sponge was observed that exhibits a dense dermal layer of short strongyles. Two hadromerid-type spicule arrangements were found. One exhibits similarities to chondrillid sponges that possess only large dermal asterose-type microsclers. The other displays a well-developed dermal layer of putative asterose microsclers and a choanosomal skeleton of randomly ornientated monaxone spicules. Few specimens of putative “Keratose” demosponge were also discovered. In combination, this paleontological evidence further corroborates a Precambrian origin of the phylum Porifera.
4. Various cuts of *Anabarella* shells were found in thin sections. Size and ornamentation give us confidence that they were correctly identified and support the stratigraphic classification of the phosphates in the late Fortunian to Stage 2.
5. The phosphatic sediment displays many different microbialite fabrics and structural types. Oncoidal-like microbialites are abundant; small domal stromatolites are less common. Both components contain significant amounts of organic matter. Long filamentous structures found resemble those of Sulfide Oxidizing bacteria (SOX-B). Because of these features, it is thought that most of the Fontanarejo apatite was formed via microbial mediation. The rapid precipitation of phosphatic cements indicates an intense flow of phosphor-rich fluids during early diagenesis and explains the relatively good preservation of fragile sponge skeletons and microbialites.

## Acknowledgements

The authors greatly acknowledge Prof. Dr. Sören Jensen, Dr. Iván Cortijo (Villuercas-Ibores-Jara UNESCO Global Geopark), Prof. Dr. Agustín Pieren, Prof. Dr. Marta Rodríguez-Martínez (Univ. Complutense Madrid),, and Dr. Javier Gonzáles Sanz (IGME, Madrid) for sharing their knowledge with us, and for their continuing support and valuable advice. For technical support we thank Dorothea Hause-Reitner, Axel Hackmann, Cornelia Hundertmark, and Yu Pei (University of Göttingen). Ellen Kappel (NY) is thanked for checking the English version of the manuscript. PSG acknowledges the support of a Postdoctoral Fellowship from the Alexander von Humboldt Foundation to stay in Göttingen. JR and JPD acknowledge the Göttingen Academy of Science and Humanities and the University of Göttingen (DFG-Courant Research Center Geobiology) for logistic and finacial support. CL thanks the National Natural Science Foundation of China (Grant No. 41972016) for financial support. This is publication number [X] of the Early Life Research Group (Department of Geobiology, Georg-August-Universität Göttingen; Göttingen Academy of Sciences and Humanities).

## References

1. Álvaro JJ, Shields-Zhou GA, Ahlberg P, Jensen S and Palacios T (2016) Ediacaran– Cambrian phosphorites from the western margins of Gondwana and Baltica. Sedimentology 63(2), 350–377.

2. Álvaro JJ, Cortijo I, Jensen S, Lorenzo S, Palacios T and Pieren AP (2019) Updated stratigraphic framework and biota of the Ediacaran and Terreneuvian in the Alcudia-Toledo Mountains of the Central Iberian Zone, Spain. Estudios Geológicos 75(2), e093.

3. Andreetto F, Dela Pierre F, Gibert L, Natalicchio M and Ferrando S (2019) Potential fossilized sulfide-oxidizing bacteria in the Upper Miocene sulfur-bearing limestones from the Lorca Basin (SE Spain): paleoenvironmental Implications. Frontiers in Microbiology 10.

4. Antcliffe JB, Callow RHT and Brasier MD (2014) Giving the early fossil record of sponges a squeeze. Biological Reviews 89(4), 972–1004.

5. Arning ET, Birgel D, Brunner B and Peckmann J (2009 a) Bacterial formation of phosphatic laminites off Peru. Geobiology 7(3), 295–307.

6. Arning ET, Lückge A, Breuer C, Gussone N, Birgel D, Peckmann J (2009 b) Genesis of phosphorite crusts off Peru. Marine Geology 262, 68–81. doi: 10.1016/j.margeo.2009.03.006

7. Arp G, Reimer A and Reitner J (2004) Microbialite formation in seawater of increased alkalinity, Satonda Crater Lake, Indonesia: reply. Journal of Sedimentary Research 74(2), 318–325.

8. Bailey JV, Corsetti FA, Greene SE, Crosby CH, Liu P and Orphan VJ (2013) Filamentous sulfur bacteria preserved in modern and ancient phosphatic sediments: implications for the role of oxygen and bacteria in phosphogenesis. Geobiology 11(5), 397–405.

9. Bandel K (1986) The reconstruction of ‘*Hyolithes kingi’* as annelid worm from the Cambrian of Jordan. Mitt. Geol.-PaIäont. Inst. 61, 35–101.

10. Birgel D, Guido A, Liu, X, Hinrichs K-U, Gier S and Peckmann J (2014) Hypersaline conditions during deposition of the Calcare di Base revealed from archaeal di- and tetraether inventories. Organic Geochemistry 77, 11–21.

11. Borchiellini C, Chombard C, Manuel M, Alivon E, Vacelet J and Boury-Esnault N (2004) Molecular phylogeny of Demospongiae: implications for classification and scenarios of character evolution. Molecular Phylogenetics and Evolution 32(3), 823–837.

12. Botting JP and Peel JS (2016) Early Cambrian sponges of the Sirius Passet Biota, North Greenland. Papers in Palaeontology 2(4), 463–487.

13. Botting JP and Muir LA (2018) Early sponge evolution: A review and phylogenetic framework. Palaeoworld 27(1), 1–29.

14. Botting JP, Muir LA and Lin JP (2013) Relationships of the Cambrian Protomonaxonida (Porifera). Palaeontologia Electronica 16(2), 9A.

15. Brain CK, Prave AR, Hoffmann, K-H, Fallic, AE, Botha A, Herd, DA, Sturrock C, Young I, Condon DJ and Allison SG (2012) The first animals: ca. 760-million-year-old sponge-like fossils from Namibia. 108(1/2), 658–665.

16. Brasier MD, Green O and Shields G (1997) Ediacarian sponge spicule clusters from southwestern Mongolia and the origins of the Cambrian fauna. Geology 25(4), 303–306.

17. Brasier MD, Perejón A and De San José (1979) Discovery of an important fossiliferous Precambrian-Cambrian sequence in Spain. Estudios Geológicos 35, 379–383.

18. Brock J (2011) Impact of sulfide-oxidizing bacteria on the phosphorus cycle in marine sediments. Bremen: Diss University of Bremen, 94p.

19. Boury-Esnault N (2002) Order Chondrisida Boury-Esnault & Lopés, 1985. Family Chondrillidae Gray, 1872. In Systema Porifera: a guide to the classification of sponges (eds. J. N. A. Hooper & R. W. M. Van Soest), pp. 52–68., New York: Kluwer Academic/Plenum Publishers.

20. Castaño Prieto RM (1987) Estudio petrológico de 23 muestras suministradas por “Minas de Almadén y Arrayanes” correspondientes a los sondeos de fosfatos de las localidades de Horcajo de los Montes, Robledo del Mazo, Fontanarejo, MAYASA company,p.

21. Chang S, Feng Q, Clausen S and Zhang L (2017) Sponge spicules from the lower Cambrian in the Yanjiahe Formation, South China: The earliest biomineralizing sponge record. Palaeogeography, Palaeoclimatology, Palaeoecology 474, 36– 44.

22. Chang S, Zhang L, Clausen S, Bottjer DJ and Feng Q (2019) The Ediacaran-Cambrian rise of siliceous sponges and development ofmodern oceanic ecosystems. *Palaeogeography, Palaeoclimatology*, Palaeoecology 333, 105438.

23. Chen J, Hou X and Lu H (1989) Lower Cambrian leptomitids (Demospongea), Chengjiang, Yunnan. Acta Palaeontologica Sinica 28(1), 17–30.

24. Chen J, Hou X and Li G (1990) New lower Cambrian demosponges -- Quadrolaminiella gen. nov. from Chengjiang, Yunnan. Acta Palaeontologica Sinica 29(4), 402– 414.

25. Clites EC, Droser ML and Gehling JG (2012) The advent of hard-part structural support among the Ediacara biota: Ediacaran harbinger of a Cambrian mode of body construction. Geology 40(4), 307–310.

26. Connes R, Carriére D and Paris J (1978) Étude du developpement des gemmules chez la demosponge marine Suberites domuncula (Olivi) Nardo. Annales des Sciences Naturelles, Zoologie, Paris 20(4), 357–387.

27. Cook PJ and Shergold JH (1984) Phosphorus, phosphorites and skeletal evolution at the Precambrian—Cambrian boundary. Nature 308, 231–236.

28. Crosby CH and Bailey JV (2012) The role of microbes in the formation of modern and ancient phosphatic mineral deposits. Frontiers in Microbiology 3(JUL).

29. Cunningham JA, Liu AG, Bengtson S and Donoghue PCJ (2016) The origin of animals: Can molecular clocks and the fossil record be reconciled? BioEssays 39(1), e201600120.

30. Díez Balda MA, Vegas R and Gonzáles Lodeiro F (1990) Structure, autochthonous sequences of Central Iberian Zone. In Pre-Mesozoic Geology of Iberia (eds. R. D. Dallmeyer & E. Martínez García), 172–188., Berlin, Heidelberg, New York, London, Paris, Tokyo, Hong Kong, Barcelona: Springer-Verlag.

31. Ding W-M and Qian Y (1988) Late Sinian to early Cambrian small shelly fossils from Yangjiaping, Shimen, Hunan. Acta Micropalaeontologica Sinica 5(1), 39–55.

32. Djerassi C and Silva CJ (1991) Biosynthetic studies of marine lipids. Accounts of Chemical Research 24(12), 371–378.

33. Dohrmann M and Wörheide G (2017) Dating early animal evolution using phylogenomic data. Scientific Reports 7(1).

34. Dohrmann M, Janussen D, Reitner J, Collins AG and Wörheide G (2008) Phylogeny and evolution of glass sponges (Porifera, Hexactinellida) (ed. F. Anderson). Systematic Biology 57(3), 388–405.

35. Ehrlich H (2010) Chitin and collagen as universal and alternative templates in biomineralization. International Geology Review 52(7–8), 661–699.

36. Ehrlich H (2019) Enigmatic structural protein spongin. In Marine Biological Materials of Invertebrate Origin (ed. H. Ehrlich), 161–172. Biologically-Inspired Systems 13, Springer.

37. Ehrlich H, Rigby JK, Botting JP, Tsurkan MV, Werner C, Schwille P, Petrášek Z, Pisera A, Simon P, Sivkov VN, Vyalikh DV, Molodtsov SL, Kurek D, Kammer M, Hunoldt S, Born R, Stawski D, Steinhof A, Bazhenov V and Geisler T (2013) Discovery of 505-million-year old chitin in the basal demosponge Vauxia gracilenta. Scientific Reports 3(3497).

38. Ehrlich H, Bazhenov VV, Debitus C, de Voogd N, Galli R, Tsurkan MV, Wysokowski M, Meissner H, Bulut E, Kaya M and Jesionowski T (2017) Isolation and identification of chitin from heavy mineralized skeleton of Suberea clavata (Verongida: Demospongiae: Porifera) marine demosponge. International Journal of Biological Macromolecules 104, 1706–1712.

39. Einsele W (1936) Über die Beziehungen des Eisenkreislaufs zum Phosphatkreislauf im eutrophen See. Archiv für Hydrobiologie 29, 664–686.

40. Ereskovsky AV (2010) The Comparative Embryology of Sponges, Dordrecht: Springer Netherlands, 323p.

41. Erpenbeck D and Wörheide G (2007) On the molecular phylogeny of sponges (Porifera). Zootaxa 1668, 107–126.

42. Erpenbeck D, Sutcliffe P, Cook S de C, Dietzel A, Maldonado M, van Soest RWM, Hooper JNA and Wörheide G (2012) Horny sponges and their affairs: On the phylogenetic relationships of keratose sponges. Molecular Phylogenetics and Evolution 63(3), 809–816.

43. Erwin DH, Laflamme M, Tweedt SM, Sperling EA, Pisani D and Peterson KJ (2011) The Cambrian Conundrum: Early divergence and later ecological success in the early history of animals. Science 334(6059), 1091–1097.

44. Finks RM and Rigby JK (2004) Paleozoic Demosponges. In Treatise on Invertebrate Paleontology, Part E Porifera, Vol.3 (eds. R. M. Finks, R. E. H. Reid, & J. K. Rigby), pp. 9–173., Boulder, Colorado and Lawrence, Kansas: Geological Society of America and University of Kansas.

45. Föllmi KB (1996) The phosphorus cycle, phosphogenesis and marine phosphate-rich deposits. Earth-Science Reviews 40(1–2), 55–124.

46. Friesenbichler E, Richoz S, Baud A, Krystyn L, Sahakyan L, Vardanyan S, Peckmann J, Reitner J and Heindel K (2018) Sponge-microbial build-ups from the lowermost Triassic Chanakhchi section in southern Armenia: Microfacies and stable carbon isotopes. Palaeogeography, Palaeoclimatology, Palaeoecology 490, 653–672.

47. Fromont J, Usher KL, Sutton DC, Toze S and Kuo J (2008) Species of the sponge genus Chondrilla (Demospongiae: Chondrosida: Chondrillidae) in Australia. Records of the Western Australian Museum 24(4), 469–486.

48. Gabaldón López V, Hernández Urroz J, Lorenzo Álvarez S, Picard Boira J, Santamaría Casanova J and Solé Pont FJ (1989) Sedimentary facies and stratigraphy of Precambrian-Cambrian phosphorites on the Valdelacasa anticline, Central Iberian Zone, Spain. In Phosphate Deposits of the World. Vol. 2. Phosphate rock resources (eds. A. J. G. Notholt, R. P. Sheldon, & D. F. Davidson), pp. 422–428., Cambridge, New York, Port Chester, Melbourne, Sydney: Cambridge University Press.

49. Gehling JG and Rigby JK (1996) Long-expected songes from the Neoproterozoic Ediacara Fauna, Pound Subgroup, South Australia. Journal of Paleontology 70, 185–195.

50. Giner JL (1993) Biosynthesis of marine sterol side chains. Chemical Reviews 93(5), 1735–1752.

51. Giner JL, Zhao H, Boyer GL, Satchwell MF and Andersen RA (2009) Sterol chemotaxonomy of marine pelagophyte algae. Chemistry & Biodiversity 6, 1111–1130.

52. Gold DA, Grabenstatter J, Mendoza A de, Riesgo A, Ruiz-Trillo I and Summons R (2016) Sterol and genomic analyses validate the sponge biomarker hypothesis. Proceedings of the National Academy of Sciences 113(10), 2684–2689.

53. Goldhammer T, Brüchert V, Ferdelman TG and Zabel M (2010) Microbial sequestration of phosphorus in anoxic upwelling sediments. Nature Geoscience 3(8), 557–561.

54. Guo J, Li Y, Han J, Zhang X, Zhang Z, Ou Q, Liu J, Shu D, Maruyama S and Komiya T (2008) Fossil association from the Lower Cambrian Yanjiahe Formation in the Yangtze Gorges area, Hubei, South China. Acta Geologica Sinica 82(6), 1124– 1132.

55. Gutiérrez-Marco JC, Lorenzo S and Sá AA (2017) Fontanarejo (Ciudad Real): una localidad icnológica excepcional del Ordovícico Inferior en los Montes de Toledo meridionales. Geogaceta 62, 47–50.

56. Hallmann C, Nettersheim BJ, Brocks JJ, Schwelm A, Hope JM, Not F and Stuhr M (2020) Reply to: Sources of C30 steroid biomarkers in Neoproterozoic-Cambrian rocks and oils. Nature Ecology and Evolution 4, 37–39.

57. Hartman WD (1980) Systematics of the Porifera. In Living and Fossil Sponges, Notes for a Short Course (eds. W. D. Hartman, J. W. Wendt, & F. Wiedenmayer), 24–51. *Sedimenta* 8, Miami: Rosenstiel School of Marine and Atmospheric Science.

58. Hofheinz W and Oesterhelt G (1979) 24-isopropylcholesterol and 22-Dehydro-24-isopropylcholesterol, novel sterols from a sponge. Helvetica Chimica Acta 62(4), 1307–1309.

59. Hooper JNA (2002) Family Hemiasterellidae Lendenfeld, 1889. In Systema Porifera: A Guide to the Classification of Sponges (eds. J. N. A. Hooper & R. W. M. Van Soest), pp. 186–195., Boston, MA: Springer.

60. Hooper JNA and Van Soest RWM. (2002) Systema Porifera: A Guide to the Classification of Sponges, New York, Boston, Dordrecht, London, Moscow: Kluwer Academic/Plenum Publishers, 1707p.

61. IGME, MAYASA AND ENCASUR 1984–1987. Proyecto de exploración sistemática coordinada de las zonas de reserva “Hespérica”, “Valdelacasa”, “Alcudia” y “Guadalupe”. Vol. 4 Fosfatos, Capítulo V, IGME.

62. Izawa MRM, Banerjee NR, Shervais JW, Flemming RL, Hetherington CJ, Muehlenbachs K, Schultz C, Das D and Hanan BB (2019) Titanite mineralization of microbial bioalteration textures in Jurassic volcanic glass, Coast Range Ophiolite, California. Frontiers in Earth Science 7.

63. Jensen S, Palacios T and Mus MM (2010) Revised biochronology of the Lower Cambrian of the Central Iberian zone, southern Iberian massif, Spain. Geological Magazine 147(5), 690–703.

64. Jones WC (1970) The composition, development, form and orientation of calcareous sponge spicules. In The Biology of the Porifera (ed. W. G. Fry), pp. 91–123. Symp. Zool. Soc. 25, London: Academic Press.

65. Jørgensen BB and Gallardo VA (1999) Thioploca spp.: filamentous sulfur bacteria with nitrate vacuoles. FEMS Microbiology Ecology 28(4), 301–313.

66. van Kempen TMG (1990) The oldest tetraxon megasclers. In New perspectives in sponge biology (ed. K. Rützler), pp. 9–16., Washington, D. C.: Smithsonian Institution Press.

67. Lévi C (1957) Ontogeny and systematics in sponges. Systematic Zoology 6, 174–183.

68. Leys SP (2003) Comparative study of spiculogenesis in demosponge and hexactinellid larvae. Microscopy Research and Technique 62(4), 300–311.

69. Leys SP, Mackie GO and Reiswig HM (2007) The biology of glass sponges (ed. B.-A. in M. Biology). Advance in Marine Biology 52, 1–147.

70. Li C-W, Chen J-Y and Hua T-E (1998) Precambrian sponges with cellular structures. Science 279(5352), 879.

71. Liñán E, Gozalo R, Palacios T, Gámez Vinaned JA, Ugidos JM and Mayoral E (2002) Cambrian. In The Geology of Spain (eds. W. Gibbons & T. Moreno), pp. 17–29., London: The Geological Society.

72. López Díaz F (1994) Estratigrafía de los materiales anteordovícicos del anticlinal de Navalpino (Zona Centroibérica). Revista de la Sociedad Geológica de España 7, 31–45.

73. López Díaz F (1995) Late Precambrian series and structures in the Navalpino Variscan Anticline (Central Iberian Peninsula). Geologische Rundschau 84, 151–163.

74. Love GD and Summons RE (2015) The molecular record of Cryogenian sponges – a response to Antcliffe (2013). Palaeontology 58, 1131–1136.

75. Love GD, Grosjean E, Stalvies C, Fike DA, Grotzinger JP, Bradley AS, Kelly AE, Bhatia M, Meredith W, Snape CE, Bowring SA, Condon DJ and Summons RE (2009) Fossil steroids record the appearance of Demospongiae during the Cryogenian period. Nature 457, 718–721.

76. Love GD, Zumberge JA, Cárdenas P, Sperling EA, Rohrssen M, Grosjean E, Grotzinger JP and Summons RE (2020) Sources of C30 steroid biomarkers in Neoproterozoic-Cambrian rocks and oils. Nature Ecology and Evolution. Nature Ecology and Evolution 4(1), 34–36.

77. Luo C and Reitner J (2014) First report of fossil “keratose” demosponges in Phanerozoic carbonates: preservation and 3-D reconstruction. Naturwissenschaften 101(6), 467–477.

78. Luo C and Reitner J (2016) ‘Stromatolites’ built by sponges and microbes - a new type of Phanerozoic bioconstruction. Lethaia 49(4), 555–570.

79. Luo C and Reitner J (2019) Three-dimensionally preserved stem-group hexactinellid sponge fossils from lower Cambrian (Stage 2) phosphorites of China. PalZ 93, 187–194.

80. Luo C, Pan B and Reitner J (2017) Chambered structures from the Ediacaran Dengying Formation, Yunnan, China: comparison with the Cryogenian analogues and their microbial interpretation. Geological Magazine 154(6), 1269–1284.

81. Luo C, Zhao F and Zeng H (2020) The first report of a vauxiid sponge from the Cambrian Chengjiang Biota. Journal of Paleontology 94(1), 28–33.

82. Maldonado M (2006) The ecology of the sponge larva. Canadian Journal of Zoology 84(2), 175–194.

83. Maldonado M (2009) Embryonic development of verongid demosponges supports the independent acquisition of spongin skeletons as an alternative to the siliceous skeleton of sponges. Biological Journal of the Linnean Society 97, 427–447.

84. Maloof AC, Rose CV, Beach R, Samuelsson BM, Calmet CC, Erwin DH, Poirier GR, Yao N and Simons FJ (2010) Possible animal-body fossils in pre-Marinoan limestones from South Australia. Nature Geoscience 3, 653–659.

85. Martínez Catalán JR, Martínez Poyatos D and Bea F (2004) Zona Centroibérica. In Geología de España (ed. J. A. Vera), pp. 68–133., IGME.

86. MAYASA, ITGE and ENCASUR 1987–1990. Exploración e investigación de fosfatos sedimentarios en las reservas “Hespérica 1 a 7” y “Malagón” y de sustancias metálicas en las reservas “Valdelacasa”, “Alcudia” y “Guadalupe”. 2a Fase (1987–1990). Tomo 2a: Exploración geológico-minera de fosfatos sedimentarios, IGME.

87. MAYASA, ITGE and ENCASUR 1990–1993 Proyecto de Investigación de la Zona Centroibérica. 3a Fase (1990–1993). Tomo 2: Exploración e investigación de fosfatos sedimentarios, IGME.

88. McCaffrey MA, Moldowan MJ, Lipton PA, Summons RE, Peters KE, Jeganathan A and Watt DS (1994) Paleoenvironmental implications of novel C30 steranes in Precambrian to Cenozoic Age petroleum and bitumen. Geochimica et Cosmochimica Acta 58(1), 529–532.

89. Mehl D (1996) Phylogenie und Evolutionsökologie der Hexactinellida (Porifera) im Paläozoikum. Geologische Paläontologische Mitteilungen der Universität Innsbruck 4, 1–55.

90. Mehl-Janussen D (1999) Die frühe Evolution der Porifera: Phylogenie und Evolutionsökologie der Poriferen im Paläozoikum mit Schwerpunkt der desmentragenden Demospongiae (‘Lithistide’). Münchner Geowiss. Abh. (A) 37, 1–72.

91. Minchin EA (1900) Chapter III. Sponges. In A treatise on zoology. Part II. The Porifera and Coelenterata (ed. E. R. Lankester), 1–178. London: Adam & Charles Black.

92. Monteserín López V, Nozal Martín F, López Díaz J, Rubio Pascual FJ and Serrano García A (1989) Mapa Geológico de España, escala 1:50.000. Hoja 735, El Robledo.

93. Morrow C and Cárdenas P (2015) Proposal for a revised classification of the Demospongiae (Porifera). Frontiers in Zoology 12(7).

94. Mortimer CH (1941) The exchange of dissolved substances between mud and water in lakes. J. Ecol. 29, 280–329.

95. Müller WEG, Li J, Schröder HC, Qiao L and Wang X (2007) The unique skeleton of siliceous sponges (Porifera; Hexactinellida and Demospongiae) that evolved rst from the Urmetazoa during the Proterozoic: a review. Biogeosciences 4, 219–232.

96. Muscente AD, Hawkins AD and Xiao S (2015) Fossil preservation through phosphatization and silicification in the Ediacaran Doushantuo Formation (South China): a comparative synthesis. Ediacaran Environments and Ecosystems 434, 46–62.

97. Nettersheim BJ, Brocks JJ, Schwelm A, Hope JM, Not F, Lomas M, Schmidt C, Schiebel R, Nowack ECM, De Deckker P, Pawlowski J, Bowser SS, Bobrovskiy I, Zonneveld K, Kucera M, Stuhr M and Hallmann C (2019) Putative sponge biomarkers in unicellular Rhizaria question an early rise of animals. Nature Ecology & Evolution 3(4), 577–581.

98. Nürnberg GK and Peters RH (1984) Biological availability of soluble reactive phosphorus in anoxic and oxic freshwater. Can. J. Fish. Aquat. Sci. 41, 757–765.

99. Peckmann J, Thiel V, Reitner J, Taviani M, Aharon P and Michaelis W (2004) A microbial mat of a large sulfur bacterium preserved in a Miocene methane-seep limestone. Geomicrobiology Journal 21(4), 247–255.

100. Peng SC, Babcock LE and Ahlberg L (2020) Chapter 19 The Cambrian Period. In Geologic Time Scale 2020 Volume 2 (eds FM Gradstein, JG Ogg, MD Schmitz and GM Ogg), pp. 565–629. Elsevier.

101. Perconig E, Vázquez Guzmán F, Velando F and Leyva F (1983) Sobre el descubrimiento de fosfatos sedimentarios en el Precámbrico Superior de España. Boletín Geológico y Minero 114, 187–207.

102. Perconig E, Vázquez Guzmán F, Velando F and Leyva F (1986) Proterozoic and Cambrian phosphorites deposits: Fontanarejo, Spain. In Phosphate Deposits of the World, Vol. 1 Proterozoic and Cambrian phosphorites (eds. P. J. Cook & J. H. Shergold), 220–234. Cambridge, London, New York: Cambridge University Press.

103. Picart Boira J (1988) Faciès et interprétation des gisements de phosphates du Cambrien inférieur de Fontanarejo, zone Centro-Ibérique (Ciudad Real). In Congreso geológico de España 2, 157–160, Granada.

104. Pieren AP and García-Hidalgo JF (1999) El Alcudiense Superior del anticlinal de Alcudia revisitado (Ciudad Real, España central). In ANNUAL MEETING OF IGCP PROJECT 376 (Laurentia-Gondwana connections before Pangea) Extended Abstracts, 207–214.

105. Pieren Pidal AP (2000) Las sucesiones anteordovícias de la region orientel de la provincial de Badajoz y área contigua de la de Ciduad Real. Doctoral dissertation. Madrid: Univ. Computense de Madrid,p.

106. Pronzato R, Pisera A and Manconi R (2017) Fossil freshwater sponges: taxonomy, geographic distribution, and critical review. Acta Palaeontologica Polonica 62(3), 467–495.

107. Redmond NE, Morrow CC, Thacker RW, Diaz MC, Boury-Esnault N, Cárdenas P, Hajdu E, Lôbo-Hajdu G, Picton BE, Pomponi SA, Kayal E and Collins AG (2013) Phylogeny and systematics of Demospongiae in light of new Small-subunit ribosomal DNA (18S) sequences. Integrative and Comparative Biology 53(3), 388–415.

108. Reid REH (1958) A monograph of the Upper Cretaceous Hexactinellida of Great Britain and Northern Ireland, part I, London, 46p.

109. Reitner J (1991) Phylogenetic aspects and new descriptions of spicule-bearing hadromerid sponges with a secondary calcareous skeleton (Tetractinomorpha, Demospongiae). In Fossil and Recent Sponges (eds. J. Reitner & H. Keupp), pp. 179–211, Berlin, Heidelberg: Springer.

110. Reitner J (1992) ‘Coralline Spongien’ Der Versuch einer phylogenetisch-taxonomischen Analyse. Berliner Geowissenschaftliche Abhandlungen (Reihe E) 1, 1–352.

111. Reitner J (1998) Early Proterozoic microbial benthic community-Basic Sponge Model: Vorstellungsbericht, Akademie der Wissenschaften Göttingen. Jahrbuch der Akademie der Wissenschaften zu Göttingen 1998, 230–235.

112. Reitner J and Neuweiler F (1995) Mud mounds: a polygenetic spectrum of fine-grained carbonate buildups. Facies 32, 1–70.

113. Reitner J, Neuweiler F and Gautret P (1995) Modern and fossil automicrites: Implications for Mud Mound genesis. In Mud mounds: a polygenetic spectrum of fine-grained carbonate buildups (eds. J. Reitner & F. Neuweiler), *Facies* 32, 4–17.

114. Reitner J and Mehl D (1995) Early Paleozoic diversification of sponges: new data and evidences. Geologisch-paläontologische Mitteilungen Innsbruck 20, 335–347.

115. Reitner J and Mehl D (1996) Monophyly of the Porifera. Verhandlungen des Naturwissenschaftlichen Vereins in Hamburg 36, 5–32.

116. Reitner J and Schumann-Kindel G (1997) Pyrite in mineralized sponge tissue - product of sulfate reducing sponge-related bacteria? Facies 36, 272–276.

117. Reitner J and Wörheide G (2002) Non-lithistid fossil Demospongiae-origins of their palaeobiodiversity and highlights in history of preservation. In Systema Porifera: a guide to the classification of sponges (eds. J. N. A. Hooper & R. W. M. Van Soest), pp. 52–68., New York: Kluwer Academic/Plenum Publishers.

118. Reitner J and Luo C (2019) Phosphorites as a new window to early Cambrian evolution of sponges: Fontanarejo, central Sapin and basal Niutitang Formation, Hunan, China. In IMECT short Abstracts, Estudios Geológicos 75, 34–36.

119. Reitner J, Luo C and Duda JP (2012) Early sponge remains from the Neoproterozoic-Cambrian phosphate deposits of the Fontanarejo area (central Spain). Journal of Guizhou University (Natural Sciences) 29 (Sup. 1), 184–186.

120. Rigby JK (1986) Sponges of the Burgess shale (Middle Cambrian), British Columbia, University of Toronto Press, 1–105p.

121. Rigby JK and Hou X-G (1995) Lower Cambrian demosponges and hexactinellid sponges from Yunnan, China. Journal of Paleontology 69(06), 1009–1019.

122. Rigby JK and Collins D (2004) Sponges of the Middle Cambrian Burgess Shale and Stephen Formations, British Columbia, Toronto: Royal Ontario Museum, 155p.

123. Rodríguez Alonso MD, Díez Balda MA, Perejón A, Pieren A, Liñan E, López Díaz F, Moreno F, Gámez Vintaned JA, González Lodeiro F, Martínez Poyatos D and Vegas R (2004) La secuencia litoestratigráfica del Neoproterozoico-Cámbrico Inferior, Dominio del Complejo Esquisto-Grauváquico. In Geología de España (ed. A. Vera), pp. 78–81., Madrid: Sociedad Geológica de España-Instituto Geológico y Minero de España.

124. Ruttenberg KC and Berner RA (1993) Authigenic apatite formation and burial in sediments from non-upwelling, continental margin environments. Geochimica et Cosmochimica Acta 57(5), 991–1007.

125. Salman V, Amann R, Girnth A.-C, Polerecky L, Bailey J, Høgslund S, Jessen G, Pantoja S and Schulz-Vogt HN (2011) A single-cell sequencing approach to the classification of large, vacuolated sulfur bacteria. Systematic and Applied Microbiology 34(4), 243–259.

126. San José MA (1984) Los materiales anteordovícicos de Anticlinal de Navalpino (provincias de Badajoz y Ciudad Real, España central). Cuadernos de Geología Ibérica 9, 81–117.

127. San José MA, Pieren AP, García-Hidalgo JF, Vilas L, Herranz P, Peláez J and Perejón A (1990) Ante-Ordovician Stratigraphy, autochthonous sequences of Central Iberian Zone. In Pre-Mesozoic Geology of Iberia (eds. R. D. Dallmeyer & E. Martínez García), pp. 147–159., Berlin, Heidelberg, New York, London, Paris, Tokyo, Hong Kong, Barcelona: Springer-Verlag.

128. Sará A (2002) Familiy Tethyidae Gray, 1848. In Systema Porifera: A Guide to the Classification of Sponges (eds. J. N. A. Hooper & R. W. M. Van Soest), pp. 245–265., Boston, MA: Springer.

129. Schmidt O (1870) Grundzüge einer Spongien-Fauna des atlantischen Gebietes, Leipzig: Wilhelm Engelmann, 88p.

130. Schulz HN and Schulz HD (2005) Large sulfur bacteria and the formation of phosphorite. Science 307, 416–418.

131. Schulze FE (1886) Über den Bau und das System der Hexactinelliden. Abhandlungen der Königlichen Akademie der Wissenschaften zu Berlin (Physikalisch-Mathematisch Classe) (1886), 1–97.

132. Schulze FE (1887) Report on the Hexactinellida collected by H.M.S. ‘Challenger’ during the years 1873–1876. Zoology 21, 1–513.

133. Serezhnikova EA and Ivantsov YA (2007) Fedomia mikhaili—A new spicule-bearing organism of sponge grade from the Vendian (Ediacaran) of the White Sea, Russia. Palaeoworld 16(4), 319–324.

134. Shimizu K, Cha J, Stucky GD and Morse DE (1998) Silicatein α: Cathepsin L-like protein in sponge biosilica. Proceedings of the National Academy of Sciences 95(11), 6234–6238.

135. Sorokin SJ, Ekins MG, Yang Q, and Cárdenas P (2019) A new deep-water Tethya (Porifera, Tethyida, Tethyidae) from the Great Australian Bight and an updated Tethyida phylogeny. European Journal of Taxonomy 529, 1–26.

136. Sperling EA, Robinson JM, Pisani D and Peterson KJ (2010) Where’s the glass? Biomarkers, molecular clocks, and microRNAs suggest a 200-Myr missing Precambrian fossil record of siliceous sponge spicules. Geobiology 8(1), 24–36.

137. Steiner M, Mehl D, Reitner J and Erdtmann B-D (1993) Oldest entirely preserved sponges and other fossils from the Lowermost Cambrian and a new facies reconstruction of the Yangtze platform (China). Berliner Geowissenschaftliche Abhandlungen E 9, 293–329.

138. Talavera C, Montero P, Martínez Poyatos D and Williams IS (2012) Ediacaran to Lower Ordovician age for rocks ascribed to the Schist–Graywacke Complex (Iberian Massif, Spain): Evidence from detrital zircon SHRIMP U–Pb geochronology. Gondwana Research 22(3), 928–942.

139. Taran M, Rad M and Alavi M (2018) Biosynthesis of TiO_2_ and ZnO nanoparticles by Halomonas elongata IBRC-M 10214 in different conditions of medium. BioImpacts 8(2), 81–89.

140. Uriz M-J, Turon X, Becerro MA and Agell G (2003) Siliceous spicules and skeleton frameworks in sponges: origin, diversity, ultrastructural patterns, and biological functions. Microscopy Research and Technique 62(4), 279–299.

141. Valladarez MI, Barba P and Ugidos JM (2002) Precambrian. In Geology of Spain (eds. W. Gibbons and T. Moreno), pp. 7–16., London: The Geological Society of London.

142. Veremeichik GN, Shkryl YN, Bulgakov VP, Shedko SV, Kozhemyako VB, Kovalchuk SN, Krasokhin VB, Zhuravlev YN and Kulchin YN (2011) Occurrence of a silicatein gene in glass sponges (Hexactinellida: Porifera). Marine Biotechnology 13(4), 810–819.

143. Vidal G, Palacios T, Gámez-Vintaned JA, Balda MAD and Grant SWF (1994) Neoproterozoic-early Cambrian geology and palaeontology of Iberia. Geological Magazine 131(6), 729–765.

144. Vidal G, Palacios T, Moczydłowska M and Gubanov AP (1999) Age constraints from small shelly fossils on the early Cambrian terminal Cadomian Phase in Iberia. GFF 121(2), 137–143.

145. Volkmer-Ribeiro C and Reitner J (1991) Renewed study of the type material of *Palaeospongilla chubutensis* Ott and Volkheimer (1972). In Fossil and Recent Sponges (eds. J. Reitner & H. Keupp), pp. 121–133., Berlin, Heidelberg: Springer-Verlag.

146. Wallace MW, Hood AvS, Woon EMS, Hoffmann K-H and Reed CP (2014) Enigmatic chambered structures in Cryogenian reefs: the oldest sponge-grade organisms? Precambrian Research 255, 109–123.

147. Wörheide G, Dohrmann M, Erpenbeck D, Larroux C, Maldonado M, Voigt O, Borchiellini C and Lavrov DV (2012) Chapter one - Deep phylogeny and evolution of sponges (phylum Porifera). *In* Advances in Marine Biology (eds. Mikel A. Becerro, Maria J. Uriz, Manuel Maldonado, & T. Xavier), pp. 1–78., Academic Press.

148. Xiao SH, Hu H, Yuan XL, Parsley RL and Cao R (2005) Articulated sponges from the Lower Cambrian Hetang Formation in southern Anhui, South China: their age and implications for the early evolution of sponges. Palaeogeography Palaeoclimatology Palaeoecology 220(1–2), 89–117.

149. Yin Z, Zhu M, Davidson EH, Bottjer DJ, Zhao F and Tafforeau P (2015) Sponge grade body fossil with cellular resolution dating 60 Myr before the Cambrian. Proceedings of the National Academy of Sciences 112(12), E1453–E1460.

150. Yochelson EL, Flower RH and Webers GF (1973) The bearing of the new Late Cambrian monoplacophoran genus Knightoconus upon the origin of the Cephalopoda. Lethaia 6(3), 275–309.

151. Zumberge JA, Love GD, Cárdenas P, Sperling EA, Gunasekera S, Rohrssen, M, Grosjean E, Grotzinger JP and Summons RE (2018) Demosponge steroid biomarker 26-methylstigmastane provides evidence for Neoproterozoic animals. Nature Ecology & Evolution 2(11), 1709–1714.

